# Diversity and stability of the gut microbiome of naked mole-rat (*Heterocephalus glaber*), the longest-lived rodent

**DOI:** 10.64898/2026.02.16.704739

**Authors:** Amir Rakhimov, Noriko Yasuda-Yoshihara, Masanori Arita, Kazuhiro Okumura, Yoshimi Kawamura, Kaori Oka, Hiroshi Mori, Yuichi Wakabayashi, Yoshifumi Baba, Hideo Baba, Kyoko Miura

## Abstract

The naked mole-rat is a subterranean rodent adapted to extreme hypoxia and low metabolic demands, with an exceptionally long lifespan relative to its small body size, while maintaining reproductive capacity. Using 16S rRNA gene sequencing of 24 samples and whole-metagenome sequencing of 11 samples from individuals up to 15 years of age, we characterized the gut microbiota and showed its complexity and distinctiveness compared with that of other rodents, including mice, squirrels, and rabbits. Although all animals were born and raised in a laboratory setting, the gut microbiota remained taxonomically stable across ages and retained key taxa previously reported in wild naked mole-rats (e.g., *Treponema* and *Desulfovibrio*). Metagenome-assembled genomes revealed the presence of archaeal methanogens and termite-gut-associated bacteria (e.g., *Methanobacteria* within Euryarchaeota and *Avelusimicrobium* within Elusimicrobiota), together with genes involved in hydrogen metabolism and archaeal methanogenesis. Compared with mice, the naked mole-rat gut microbiota was enriched in carbohydrate-active enzymes targeting plant cell-wall polysaccharides, resembling those found in ruminants. We also detected evidence of flagellates, ciliates, and fungi, which may further contribute to polysaccharide degradation and fermentation, potentially within the enlarged cecum. Together, this comprehensive analysis provides distinctive gut microbial features of the naked mole-rat that may be associated with the naked mole-rat’s low metabolic rate and exceptional longevity.

## Introduction

The composition of the gut microbiota is shaped not only by diet but also by host aging (Bana and Cabreiro 2019; López-Otín et al. 2023). During aging, some microorganisms change in their abundance, other species disappear entirely, or new species colonize the gut (Claesson et al. 2011; Zhao et al. 2019; Olsson et al. 2022; Borsom et al. 2023). In aged humans and mice, an increased relative abundance of Bacteroidota compared with Bacillota and an increase of opportunistic pathogens are often observed.

In the field of aging research, the longest-lived rodent, the naked mole-rat (*Heterocephalus glaber* or NMR), is an emerging biomedical model due to its unique physiology and longevity. NMRs live underground in Northeastern Africa and exhibit several distinctive traits, such as eusociality, a subterranean lifestyle, an exceptionally long lifespan (approximately 40 years), resistance to cancer, and hypoxia tolerance (Wong et al. 2022; Oka et al. 2023). NMRs display a negligible senescence phenotype (Ruby et al. 2018; Ruby et al. 2024), in which physiological functions such as fecundity, cardiac function, and body composition are maintained until late in their lives, and the mortality rate does not increase with age (Can et al. 2022; Brieño-Enríquez et al. 2023). These fascinating characteristics make NMRs an intriguing model for aging research (Buffenstein 2008; Gorbunova et al. 2014; Oka et al. 2023). More recently, their close relative, the Damaraland mole-rat (DMR), which inhabits subterranean environments in southern Africa, has also attracted increasing attention due to shared characteristics with NMRs, including eusociality, extended lifespan (maximum lifespan 20 years), and hypoxia tolerance (Yap et al. 2022).

NMRs have a unique digestive system adapted to their ecological niche. They live in underground burrows and feed primarily on tubers and plant roots, a diet low in protein. As in other rodents, fiber digestion occurs in the caecum via microbial fermentation (McBee 1977; Hume 1997). In ruminants, fermentation is exothermic, and the temperature of the gastric rumen is typically 2–3°C higher than their core body temperature. In contrast, the caecal temperature of NMRs is known to be several degrees lower (28.9 ± 1.3°C) than their core body temperature (30.2 ± 1.2°C) (Yahav and Buffenstein 1992; Buffenstein 1996). This unusually low caecal temperature suggests that the caecum may function as an internal heat sink, potentially contributing to thermal regulation within semi-enclosed burrows with high humidity. Despite the low temperature, caecum function remains efficient, with fiber digestibility reaching 90–95% (Buffenstein and Yahav 1991; Yahav and Buffenstein 1992; Buffenstein 1996). It has been suggested that the caecal microbiota of NMRs may include nitrogen-fixing bacteria, which could supply nitrogen to the microbiota and ultimately to the host (Dyer 1998). However, clear data supporting nitrogen fixation in the NMR caecum remains lacking.

NMRs also exhibit extensive coprophagic behavior, i.e., consumption of their own feces and those of colony members. They produce two types of feces: soft primary feces, which are consumed immediately or fed to young individuals or the queen, and hard secondary feces, which are rarely consumed (Dyer 1998). The soft primary feces are analogous to regurgitated cud in ruminants, which is re-chewed to facilitate digestion. These unique dietary habits further suggest a distinctive role for their gut microbiota in supporting healthy longevity in NMRs.

Until recently, the gut microbial composition of NMRs had been poorly characterized. Using plate-based culturing methods, Debebe *et al*. identified Firmicutes (currently Bacillota) and Bacteroidetes (currently Bacteroidota) as the major bacterial phyla in the NMR gut. At the species level, dominant taxa included *Bacillus megaterium*, *Bacteroides thetaiotaomicron*, *Bacteroides ovatus*, *Staphylococcus sciuri*, and *Paenibacillus spp*. (Debebe et al. 2016). In their subsequent work using 16S rRNA gene sequencing, the gut microbial composition of NMRs was shown to contain a high abundance of Spirochetaceae and Prevotellaceae, as in the microbiota of the Hadza, a native human hunter-gatherer population (Debebe et al. 2017). Another study compared the cecal and tracheal microbiota of NMRs housed in sterile or non-sterile cages, and concluded that the breeding environment has a substantial impact on microbial composition (Cong et al. 2018). More recently, Chua et al. characterized the microbiome of fecal samples collected from the toilet chamber of an NMR colony, and suggested that captivity contributes to the microbial differences between wild and laboratory-bred NMRs (Chua et al. 2022). Collectively, these studies highlight the distinctiveness and plasticity of the NMR gut microbiota; however, they remain largely limited to taxonomic descriptions based on culturing or 16S rRNA gene sequencing. As a result, the functional potential of the microbiota, its genomic diversity, and the contribution of non-bacterial microorganisms remain poorly understood, particularly in a comparative evolutionary context.

Here we comprehensively characterize the gut microbiota of laboratory-bred NMRs using both 16S rRNA gene sequencing and shotgun metagenomic sequencing. To place the NMR microbiota in an evolutionary and ecological framework, we compare it with those of 6 other species: Damaraland mole-rat in Hystricomorpha (DMR, *Fukomys damarensis*), 3 species in Myomorpha (Lesser blind mole-rat, *Nannospalax leucodon*; Siberian flying squirrel, *Pteromys volans orii*; and Laboratory mouse, *Mus musculus*), and 2 species in Lagomorpha (European brown hare, *Lepus europaeus*; and European rabbit, *Oryctolagus cuniculus*). We further reconstruct metagenome-assembled genomes (MAGs) to assess the metabolic potential of the microbiota and to identify characteristic microbial lineages associated with NMRs. Notably, our metagenomic analyses revealed the presence of diverse non-bacterial organisms, including archaea, fungi, and protozoa, which have been largely overlooked in previous studies. Functional annotation of a non-redundant gene catalog of the microbiome suggests a marked enrichment of carbohydrate-active enzymes (CAZymes) involved in the degradation of complex polysaccharides, consistent with the ability of NMRs to efficiently digest plant-derived diets. Importantly, the taxon-specific distribution of certain CAZyme subclasses suggests a tiered organization of digestive functions among distinct microbial groups. Together, this high-resolution taxonomic and functional dataset provides a valuable resource for elucidating the functional potential of NMR gut microbiome and its potential contribution to host physiology.

## Materials and Methods

### 1. Sample collection and storage

The Ethics Committees of Kumamoto University (approval no. A2022-079) and Chiba University (approval no. 23-5) approved all procedures in accordance with the Guide for the Care and Use of Laboratory Animals (United States National Institutes of Health, Bethesda, MD).

Naked mole-rat (NMR) colonies at Kumamoto University were maintained at 30°C ± 0.5°C and 55% ± 5% humidity with 12 h light and 12 h dark cycles. To collect fresh feces, male and female NMRs (n = 24), MSM/Ms mice (n = 8, all males), and FVB/N mice (n = 3, all males) were placed in disposable cages. C57BL/6 mice (n = 4) were placed in autoclaved cages. Immediately after defecation, samples were quickly immersed in liquid nitrogen and stored at −80°C. This collection method was repeated two to three times for each individual.

Sampling for whole metagenome sequencing was done a year after sampling for 16S rRNA gene sequencing. Feces from H3, H4, H15, H21, G14, and G18 were used for both 16S and whole metagenome sequencing.

### 2. 16S rRNA gene sequencing

#### 2.1. Data Acquisition

DNA extraction from the fecal samples and 16S rRNA gene amplicon sequencing were performed by Noster Corporation (Kyoto, Japan) using the QIAamp DNA Microbiome Kit. The V3–V4 region of the 16S rRNA gene was amplified using the primer pair 341F (5’-CCTACGGGNGGCWGCAG-3’) and 805R (5’-GACTACHVGGGTATCTAATCC-3’) (Claesson et al. 2010). For the 150 bp paired-end sequencing with the MiSeq system (Illumina), the MiSeq Reagent Kit v3 and PhiX Control Kit v3 (Illumina) were used.

Fecal samples data from NMRs (n = 24) and C57BL/6 mice (B6 mouse, n = 4) were obtained at Kumamoto University. Original data from MSM/Ms mice (MSM mouse; n = 8) and FVB/N mice (FVBN mouse; n = 3) were obtained at the Chiba Cancer Center Research Institute. Public sequencing data were downloaded for DMRs (n = 20) (Bensch et al. 2022), European brown hares (*L. europaeus*; n = 9) and European rabbits (*O. cuniculus*; n = 12) (Shanmuganandam et al. 2020), Lesser blind mole-rat (*N. leucodon*; n = 15) (Sibai et al. 2020), and Siberian flying squirrel (*Pteromys volans orii*, PVO; n = 10) (Liu et al. 2020). All rRNA gene sequence data compared in this study targeted the V3-V4 region of the 16S rRNA gene (341F -805R).

#### 2.2. Data analysis

All 16S rRNA gene sequence data were analyzed using the same method with QIIME2 v2024.2 (Bolyen et al. 2019). Only forward reads were used to standardize all data conditions because the reads from *N. leucodon* (public data) did not overlap: the maximum length of the forward and reverse reads in that host were 234 nt and 231 nt, respectively, whereas the expected length of the V3-V4 amplicon length is approximately 465 nt. Because the forward and reverse reads did not overlap, paired-end read merging was not possible, precluding the use of both reads.

We excluded 1 hare and 5 rabbit samples because their hosts were pregnant and/or lactating (MF-157, MF-138, MF-141, MF-142, MF-145, MF-146).

After trimming with the q2-cutadapt plugin, reads were denoised, dereplicated, and filtered for chimeras using q2-DADA2 v2024.2.0 (Callahan et al. 2016). The processed reads were truncated to 234 nt and filtered by length (>199 nt) to remove very short reads.

The resulting reads (also known as amplicon sequence variants, or ASVs) were classified using a Naive Bayes classifier trained on the SILVA 138.1 database (Pruesse et al. 2007; Quast et al. 2013). After pre-processing with the q2-RESCRIPt plugin (`get-silva-data`, `reverse-transcribe`, `cull-seqs` commands) (Ii et al. 2021), the database was reduced to sequences containing V3-V4 primers and truncated to 234 nt (`q2-feature-classifier extract-reads` command). The truncation length was chosen to improve classification accuracy. Next, reference sequences were de-replicated in a unique mode (q2-RESCRIPt `dereplicate` command), and the classifier was trained (`q2-feature-classifier fit-classifier-naive-bayes` command). After taxonomic classification, ASVs from chloroplasts, mitochondria, and archaea were removed. The filtered ASV sequences were aligned to a tree for downstream analyses (`q2-phylogeny align-to-tree-mafft-fasttree` command).

The table of ASV abundances in each sample (ASV table), sample metadata, table of identified taxa (taxonomy table), and rooted phylogenetic tree were imported into RStudio v2023.12.1+402 of R v4.4.3 (2025-02-28 ucrt) (RStudio Team 2020; R Core Team 2018) and converted into a phyloseq object (qiime2R `qza_to_phyloseq` function) (McMurdie and Holmes 2013; Bisanz 2023). A phyloseq object is a collection of heterogeneous data types from a microbiome study that usually consists of an ASV table, a table of sample metadata, a taxonomy table, and a phylogenetic tree. The Phyloseq package facilitates reproducible interactive analysis of microbiome data generated with different methodologies.

Nine samples with a total abundance of less than 20,000 reads were excluded (PVO_01, PVO_02, PVO_03, PVO_04, PVO_05, PVO_06, PVO_11, PVO_12, PVO_13), because they contained 10–20-fold fewer reads than other samples. These samples were excluded from all downstream analyses. The phyloseq object was transformed into three tables of taxonomic abundances corresponding to three taxonomic ranks: phylum, family, and genus (`tax_glom` function). During the transformation, only reads from the bacterial domain were retained (`subset_taxa` function).

### 3. Alpha and Beta diversity

Before calculating the Alpha diversity metrics, the genus-level table of absolute abundances was standardized for the sequencing depth in the R vegan package v2.6-4 (`rrarefy` function) with the subsampling depth equal to the smallest sample size of n = 21,384 (Roswell et al. 2021; Oksanen et al. 2022). With the rarefied table, the number of genera (`sobs` function), Shannon diversity index (`diversity` function, index = “`shannon`”), and inverse Simpson index (`diversity` function, index = “`invsimpson`”) were calculated. The Kruskal-Wallis rank sum test in each Alpha-diversity metric across animal hosts and the pairwise Wilcoxon rank sum test with Benjamini-Hochberg correction were performed in the R stats package (`kruskal.test` and `pairwise.wilcox.test` function, respectively).

Beta-diversity analysis was performed on the rarefied genus-level table of abundances using principal component analysis (PCA). The table was standardized using the R vegan package (`decostand` function, `method= “rclr”`, `logbase = “exp”`). The rclr standardization was chosen due to the compositionality of microbiome datasets and the need to perform a centered log-ratio transformation (Aitchison et al. 2000; Gloor et al. 2017; Martino et al. 2019). Finally, the principal components were calculated using R (`prcomp` function).

### 4. Differential microbial abundance analysis

To identify significantly more or less abundant bacterial genera between animal hosts, three methods were used: MaAsLin2 v1.20.0 (Mallick et al. 2021), ALDEx2 v1.38.0 (Fernandes et al. 2013; Fernandes et al. 2014; Gloor et al. 2016), and ANCOM-BC v2.8.1 (Lin and Peddada 2020). All tools used a genus-level rarefied abundance table as their input and the naked mole-rat as the reference host (Supplementary methods).

The MaAsLin2 parameters were the same as when we compared rodent hosts. In the discriminant analysis, NMR samples were divided into two age groups: young (0–9 years old, n = 18) and old (10–15 years old, n = 6) (Supplementary Table S1). We also compared males (n = 9) and females (n = 15). MaAsLin2 allows for adding a control variable by specifying which NMR samples belong to the same group (e.g., to the same individual in a longitudinal study or family in a lineage study) using the “`random_effects`” parameter.

### 5. Whole metagenome sequencing

For the whole metagenome sequencing, 11 NMRs were selected based on the 16S rRNA gene sequencing results: two different age groups, ranging from 2 to 16 years old, with exemplary microbiota. DNA was extracted with the QIAmp Fast DNA Stool Mini Kit (Qiagen), DNA concentration was measured using NanoDrop, and 150 bp pair-end sequencing was performed with Illumina HiSeq 2500 by Novogene (Japan).

Before taxonomic classification, DNA reads were decontaminated with the Bowtie2 pipeline. First, Cutadapt v4.9 was used to remove reads containing adapters, or containing N > 10%, or containing low-quality (Qscore ≤ 5) bases for over 50% of the total bases (Martin 2011). Then, reads were aligned to five reference genomes with Bowtie2 v2.5.3 (Langmead et al. 2009; Langmead and Salzberg 2012; Langmead et al. 2019): naked mole-rat (Heter_glaber.v1.7_hic_pac), human (T2T-CHM13v2.0), Damaraland mole-rat (DMR_v1.0_HiC), mouse (GRCm39), and human decoy genome (ftp://ftp.1000genomes.ebi.ac.uk/vol1/ftp/technical/reference/phase2_reference_assembly_sequence/hs37d5ss.fa.gz).

Taxonomic classification of metagenomic reads was performed with Kraken2 with a k-mer length of 30, a minimizer length of 26, a minimizer space of 6, and a confidence threshold of 0.1 (Wood et al. 2019). Species abundances were re-estimated using Bracken software with a threshold of 10 and a read length of 150 (Lu et al. 2017). The reference database for Kraken2 included bacteria, archaea, viruses, human genome, UniVec_Core, fungi, plants, protozoa, and plasmids.

### 6. Metagenome-assembled genome (MAG) assembly

Decontaminated reads were assembled into contigs with MEGAHIT v1.2.9 using default parameters (Li et al. 2015). Reads were aligned to contigs with bbwrap from BBMap v39.01 (Bushnell et al. 2017). The sorted BAM file where either read is mapped to contigs was used for the generation of depth files with jgi_summarize_bam_contig_depths and binning with MetaBAT2 v2.17 (Kang et al. 2019). Assembled MAGs were filtered with CheckM2 v1.1.0 (>= 90% completeness, <= 5% contamination), and 319 high-quality bins remained (Chklovski et al. 2023). These bins were aggregated and dereplicated with dRep v3.6.2 using a secondary clustering threshold of 95% average nucleotide identity (ANI) and at least 25% overlap between genomes (-pa 0.95, -sa 0.95, -comp 80, -con 10, -strW 0, -nc 0.25, -cm larger, -d) (Olm et al. 2017). GTDB-Tk v2.4.0 was used to assign taxonomy to representative MAGs (Chaumeil et al. 2020), and BLASTN v2.16.0 against SILVA 138.2 SSU Ref NR99 database was used to assign taxonomy to all contigs from MEGAHIT (sequence identity > 90%, alignment length > 500 bp, sorted by bitscore and extracted top hits) (Altschul et al. 1997; Camacho et al. 2009).

Genes were predicted with PROKKA v1.14.6 using the “`--metagenome`” parameter (Seemann 2014). All predicted protein sequences were aggregated, filtered by length (>100 bp, i.e., 33 amino acids), and clustered with MMseqs2 (v16.747c6) “`easy-cluster`” (--min-seq-id 0.95, -c 0.9, --cov-mode 0) (Steinegger and Söding 2017). Non-redundant protein sequences were annotated by KofamScan v1.3.0 (Aramaki et al. 2020) to predict KEGG Orthology (KO), and a standalone run_dbcan v4.1.4 to predict Carbohydrate-Active enZYmes (CAZymes) (Zheng et al. 2023). KO predictions where the threshold was empty or the score was less than the threshold were removed, then KOs were sorted by score and evalue, and the top hit was retained. To identify the putative taxonomy of CAZymes, the protein sequences were mapped to the CAZy database using BLASTP (Drula et al. 2022). Reference CAZymes sequences were downloaded from https://bcb.unl.edu/dbCAN2/download/CAZyDB.07142024.fa, and the taxonomy information was downloaded from https://www.cazy.org/IMG/cazy_data/cazy_data.zip.

Duplicate reads were removed from sorted BAM files with Picard `MarkDuplicates` v3.4.0 (Broad Institute 2019), and gene abundances were quantified using `htseq-count` from HTSeq v2.0.5 (Putri et al. 2022).

We predicted the gut temperature using the Metagenomic Thermometer (MetaThermo) developed by Kurokawa et al (Kurokawa et al. 2023). The input data were amino acid sequences of genes predicted by PROKKA.

## Results

### Microbiota of mole-rats is more diverse than in other rodents

We first compared the composition of gut microbiota between NMR and other related animals. From our own data acquisition and the data from public repositories, 16S rRNA gene sequencing results were compared for 9 animal species and strains: naked mole-rat (NMR, *H. glaber*; n = 24), Damaraland mole-rat (DMR, *F. damarensis*; n = 20), lesser blind mole-rat (*N. leucodon*; n = 15), Siberian flying squirrel (*P. volans orii*; n = 10), C57BL/6 mouse (B6, *M. musculus*; n = 4), MSM/Ms mouse (*M. musculus*; n = 8), FVB/N mouse (*M. musculus*, n = 3), European brown hare (*L. europaeus*, n = 8), and European rabbit (*O. cuniculus*, n = 7) using the same data processing pipeline. The resulting ASVs and genera per host are summarized in Table 1.

**Table 1.** Summary statistics for the final QIIME2 output based on unrarefied data.

|  | Total reads | Library size<br>± SD | ASV | Phyla | Families | Genera<br>(classified :<br>unclassified) |
| --- | --- | --- | --- | --- | --- | --- |
| Damaraland mole-rat (DMR) (n=20) | 3,226,720 | 161,336<br>± 68,602 | 1,826 | 21 | 156 | 257<br>(183 : 74) |
| Naked mole-rat (NMR) (n=24) | 1,844,935 | 76,872<br>± 26,287 | 1,848 | 19 | 111 | 231<br>(179 : 52) |
| European rabbit (n=7) | 2,417,378 | 345,340<br>± 70,861 | 3,456 | 13 | 101 | 188<br>(136 : 52) |
| Siberian flying squirrel (n=10) | 758,468 | 75,847<br>± 55,176 | 1,490 | 13 | 96 | 186<br>(149 : 37) |
| European brown hare (n=8) | 2,923,880 | 365,485<br>± 48,701 | 1,478 | 14 | 87 | 173<br>(121 : 52) |
| MSM/Ms mouse<br>(n=8) | 613,902 | 76,738<br>± 8,144 | 907 | 11 | 62 | 134<br>(110 : 24) |
| C57BL/6 mouse<br>(n=4) | 350,539 | 87,635<br>± 9,606 | 943 | 12 | 56 | 125<br>(105 : 20) |
| FVB/N mouse<br>(n=3) | 236,930 | 78,977<br>± 9,116 | 665 | 11 | 52 | 117<br>(96 : 21) |
| Lesser blind<br>mole-rat (n=15) | 539,095 | 35,940<br>± 11,383 | 1,290 | 10 | 53 | 114<br>(93 : 21) |
| **TOTAL** |  |  |  | **25** | **222** | **524** |

The ASVs were grouped into 25 phyla, 222 families, and 524 genera (Supplementary Tables S2, S3; Supplementary figure S1). When the library sizes were not corrected, the DMR samples contained the highest number of genera (257), followed by NMR (231), European rabbit (188), and Siberian flying squirrel (186). The distribution of dominant taxa did not differ much from the original reports for the public data, although only forward reads were used in our analysis (Supplementary Tables S2, S4, S5, S6). When the library sizes were standardized with the rarefy command (see Alpha and Beta diversity in Methods), the NMR samples showed the highest richness of microbial communities (Table 2 and Figure 2a, Supplementary Table S7).

**Figure 1.**
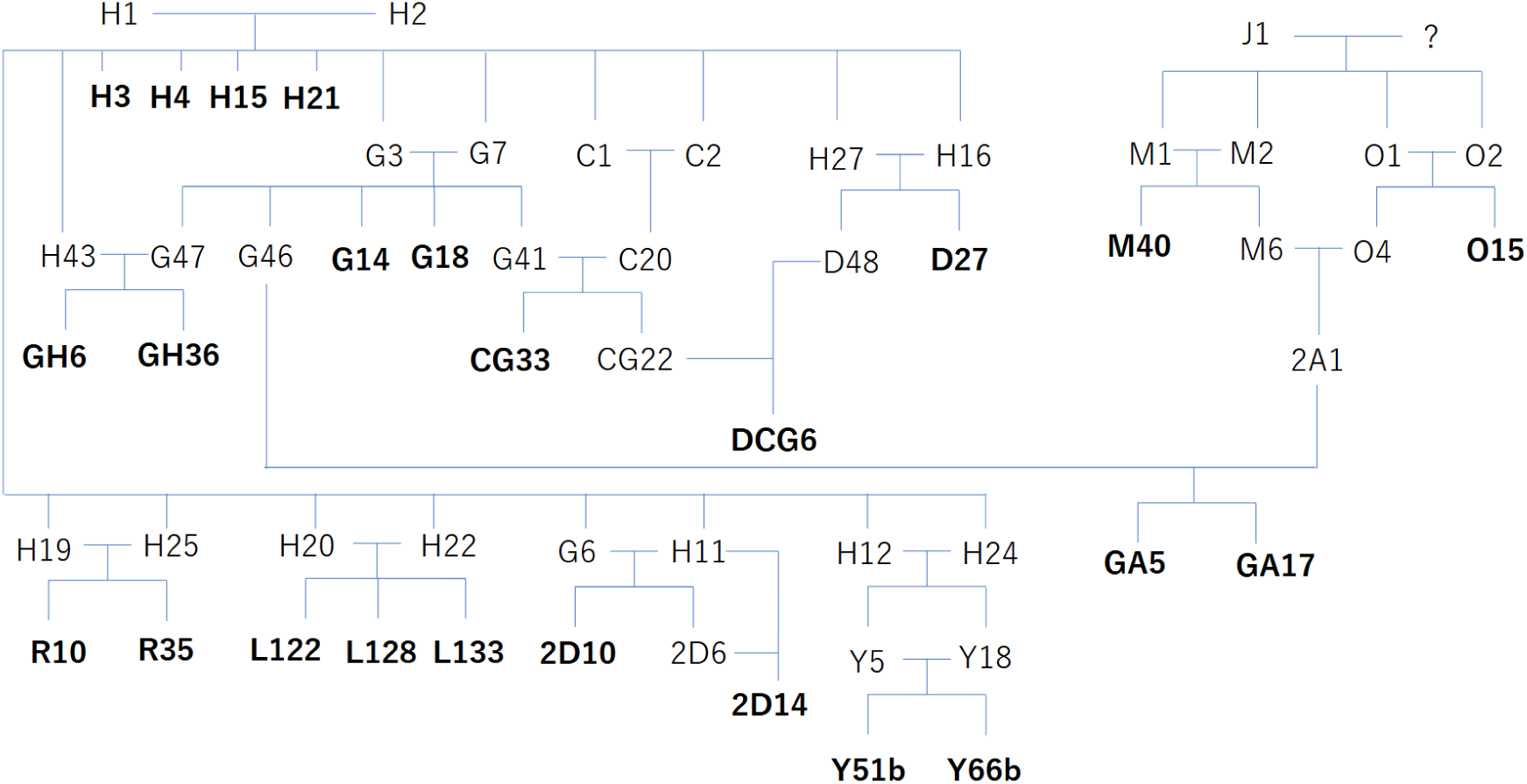
Family tree of NMR samples. Individuals used in this study are marked in bold. Most individuals are kept as a pair (king and queen) in a private cage. From larger colonies, no queen was selected.

**Figure 2.**
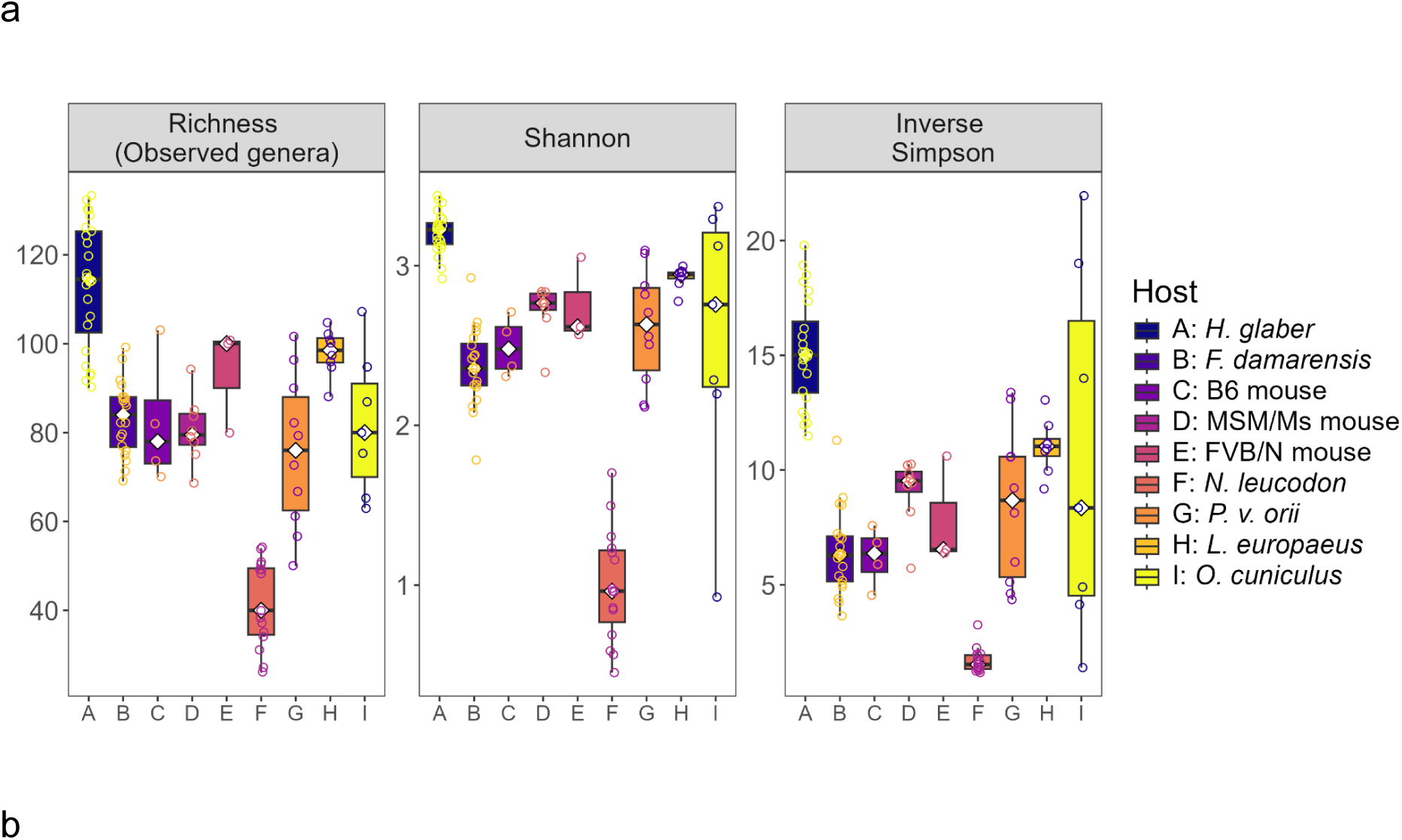
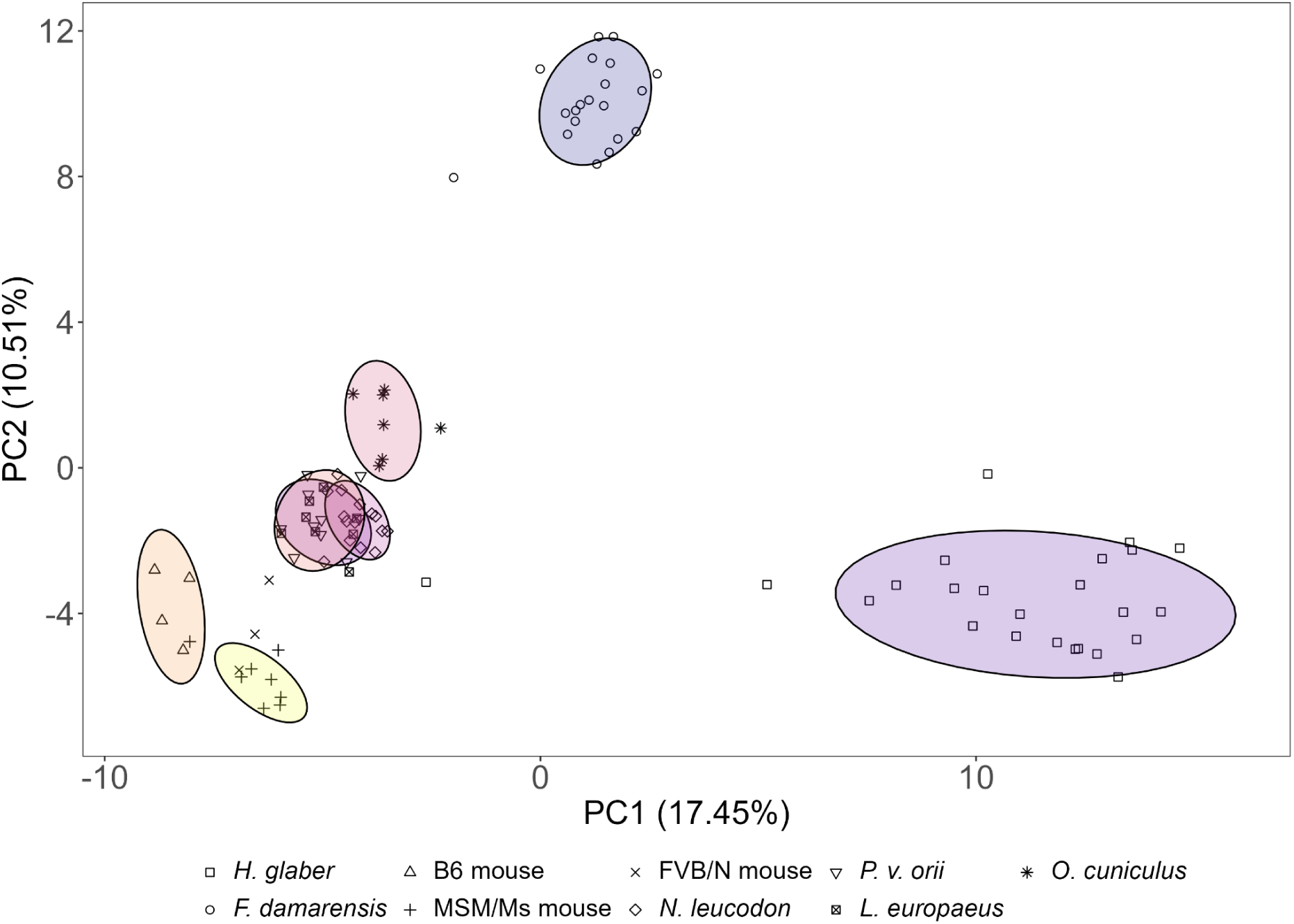
Alpha diversity of the gut microbiota and the PCA result of different rodents. **a.** Gut microbiota of naked mole-rat (leftmost plot) was the most diverse among the compared rodents in 3 Alpha-diversity indices. Dots show the individual samples. White diamonds show the medians **b.** PCA results of gut microbiota from different rodents.

**Table 2.**
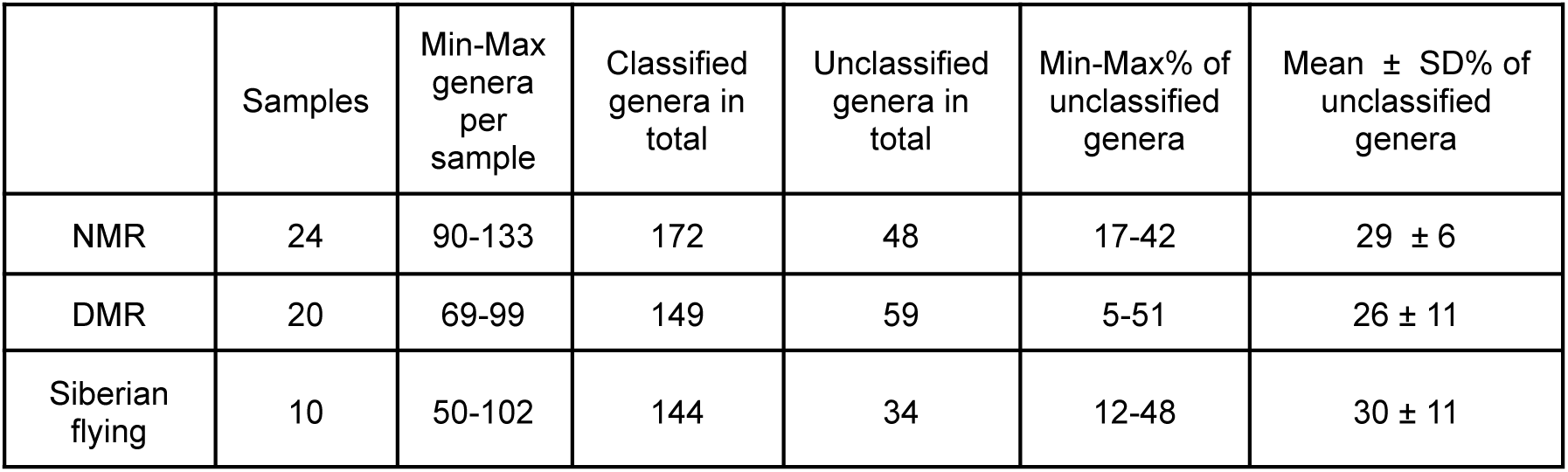
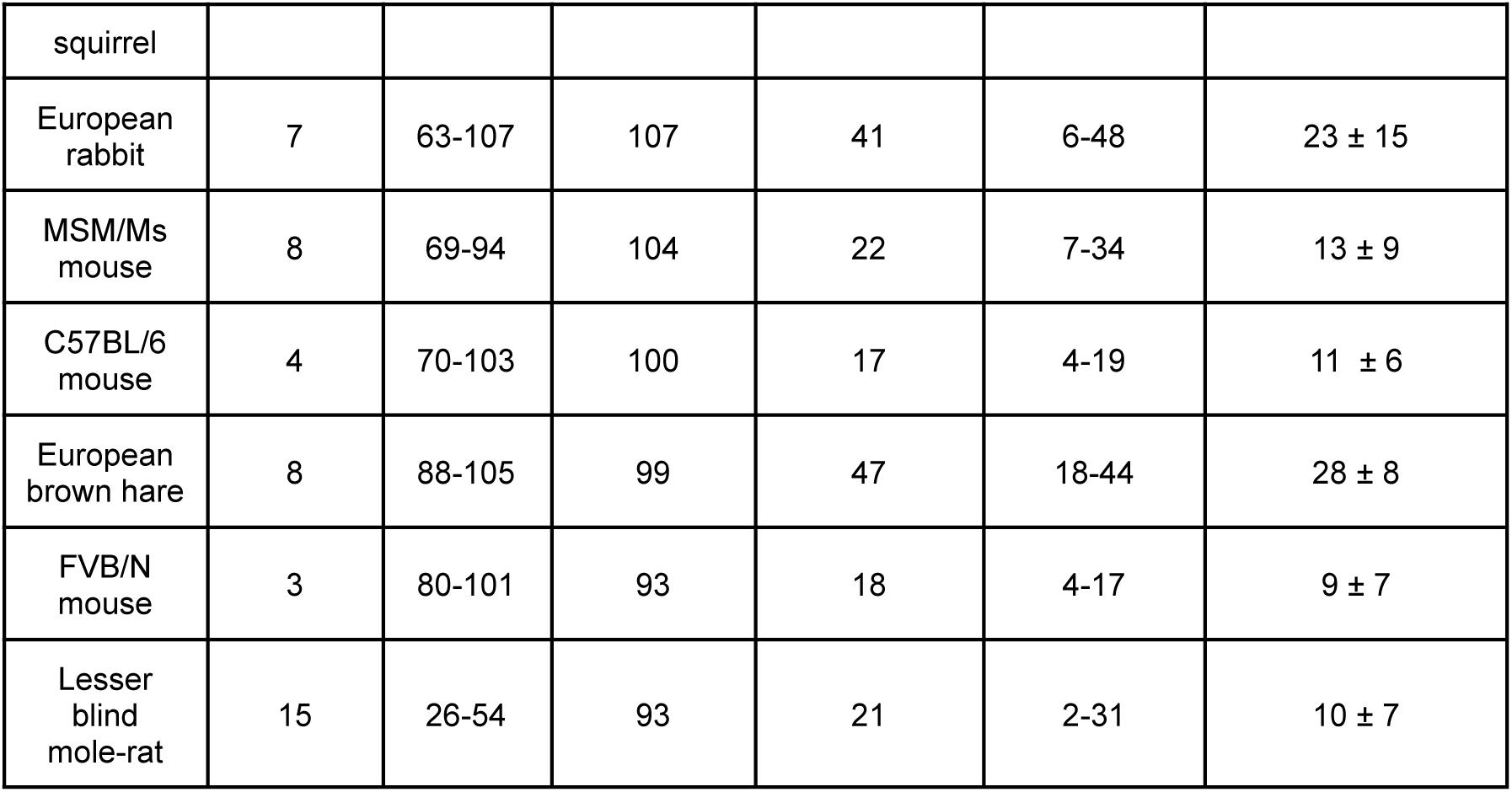
Relative abundance of classified (and unclassified) genera based on rarefied data.

As for the ratio of ‘unclassified’ genera, the Siberian flying squirrel was the highest (average 30% ± 11%), followed by NMR (average 29% ± 6%), European brown hare (average 28% ± 8%), and DMR (average 26% ± 11%) after the library-size correction. For Myomorpha species, the ratios were notably lower (Table 2).

The uniqueness of NMR and DMR was also evident in the principal component analysis (PCA) (Figure 2b). The first principal component in Figure 2b, separating NMR from other rodents, was represented by unclassified Erysipelotrichaceae genus (average 5.53% in NMR, 0.07% in DMR), *Prevotella* (3.81%, 14.2%), Prevotellaceae *UCG-003* (5.48%, 0.151%), *p-251-o5* (6.32%, 0.09%), unclassified Prevotellaceae genus (5.23%, 9.26%), and *Fibrobacter* (5.71%, absent in DMR) (Supplementary Table S10). The second principal component, separating NMR from DMR, was represented by unclassified Rikenellaceae genus (0.02%, 4.28%), unclassified Bacteroidia order (0.2%, 4.99%), *Prevotella*, unclassified Eggerthellaceae genus (0.008%, 0.45%), unclassified Gastranaerophilales (0.92%, 2.48%), and Prevotellaceae *UCG-001* (4.63%, absent in DMR).

Differential abundance analyses to identify NMR-specific bacteria showed inconsistent results among the three software programs: with MaAsLin2 and ANCOM-BC, 33 and 43 genera were found statistically abundant only in NMRs, respectively, while no difference was found with ALDEx2. Major genera found with MaAsLin2 and ANCOM-BC were *p-251-o5*, *Fibrobacter*, unclassified Erysipelotrichaceae genus, Prevotellaceae *UCG-003*, *Treponema* (2.2% in NMR), and unclassified Paludibacteraceae genus (2.17%). These names were consistent with those from the PCA (Supplementary figure S2, Supplementary Table S11).

### Age or sex does not alter the microbial composition of NMRs

Among NMR individuals, PCA did not show a noticeable relationship between age and microbial composition (Figure 3b), although the Alpha diversity decreased with age (Figure 3c). The young and the old groups had 1745 ASVs (229 genera) and 771 ASVs (157 genera), respectively, of which 668 ASVs (155 genera) were shared. Based on the number of ASVs for each taxonomic group, Lachnospiraceae (55 ASVs, 1.9% of all reads), Muribaculaceae (54 ASVs, 10.4%), and *Treponema* (42 ASVs, 1.78%), ranked high. In terms of abundance, however, 30-40% of the samples were dominated by less than 10 stable taxonomic groups: *Allobaculum* (Erysipelotrichaceae), another Erysipelotrichaceae, Eubacteriaceae genera (2 ASVs), *Fibrobacter*, *p-251-o5*, Muribaculaceae (2 ASVs), Paludibacteraceae genus, and Prevotellaceae genera (4 ASVs) (Figure 3a). These groups did not show age- or sex-dependent clusters (Figure 3b and Supplementary figure S3). The M40 sample showed anomalous composition and was excluded from the later analyses.

**Figure 3.**
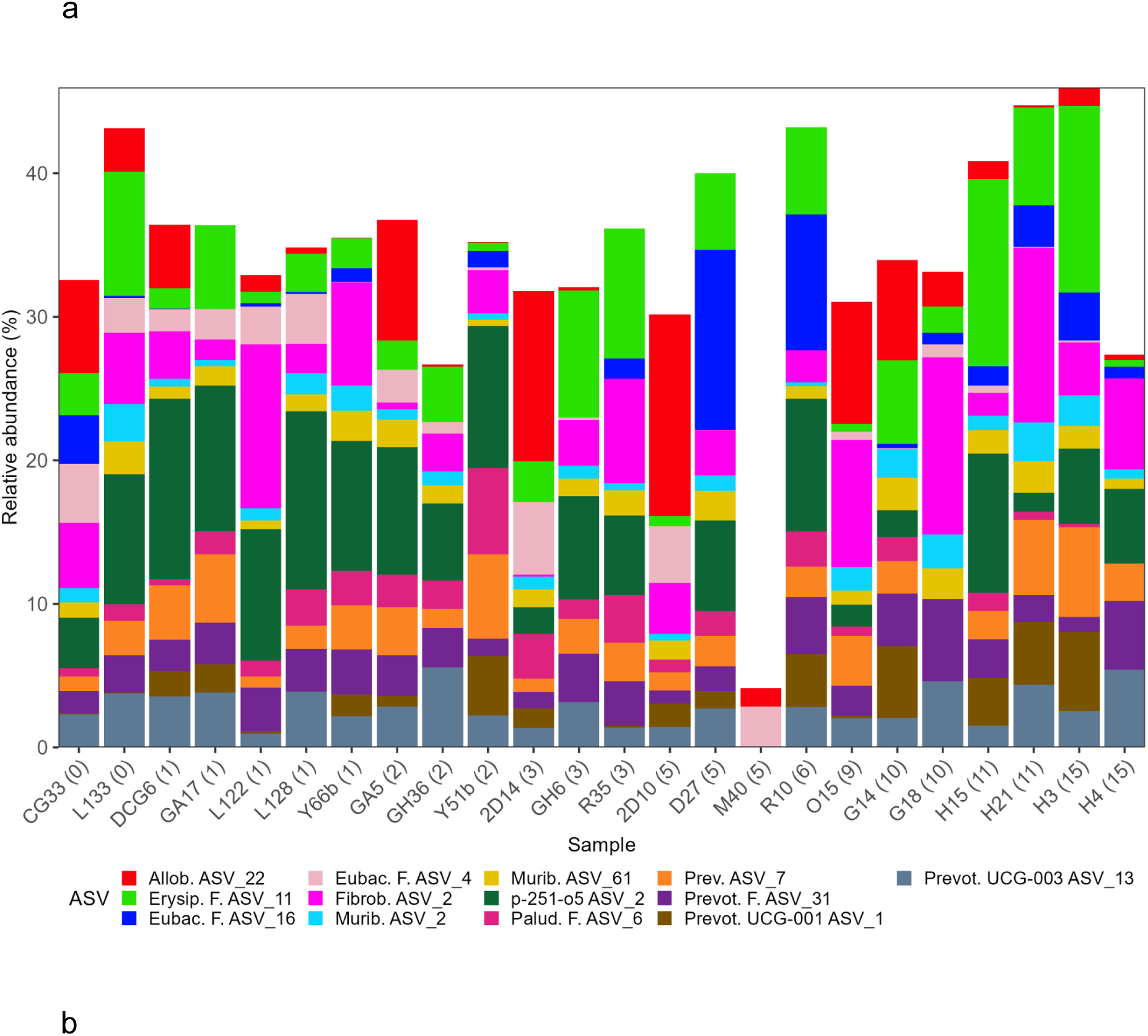
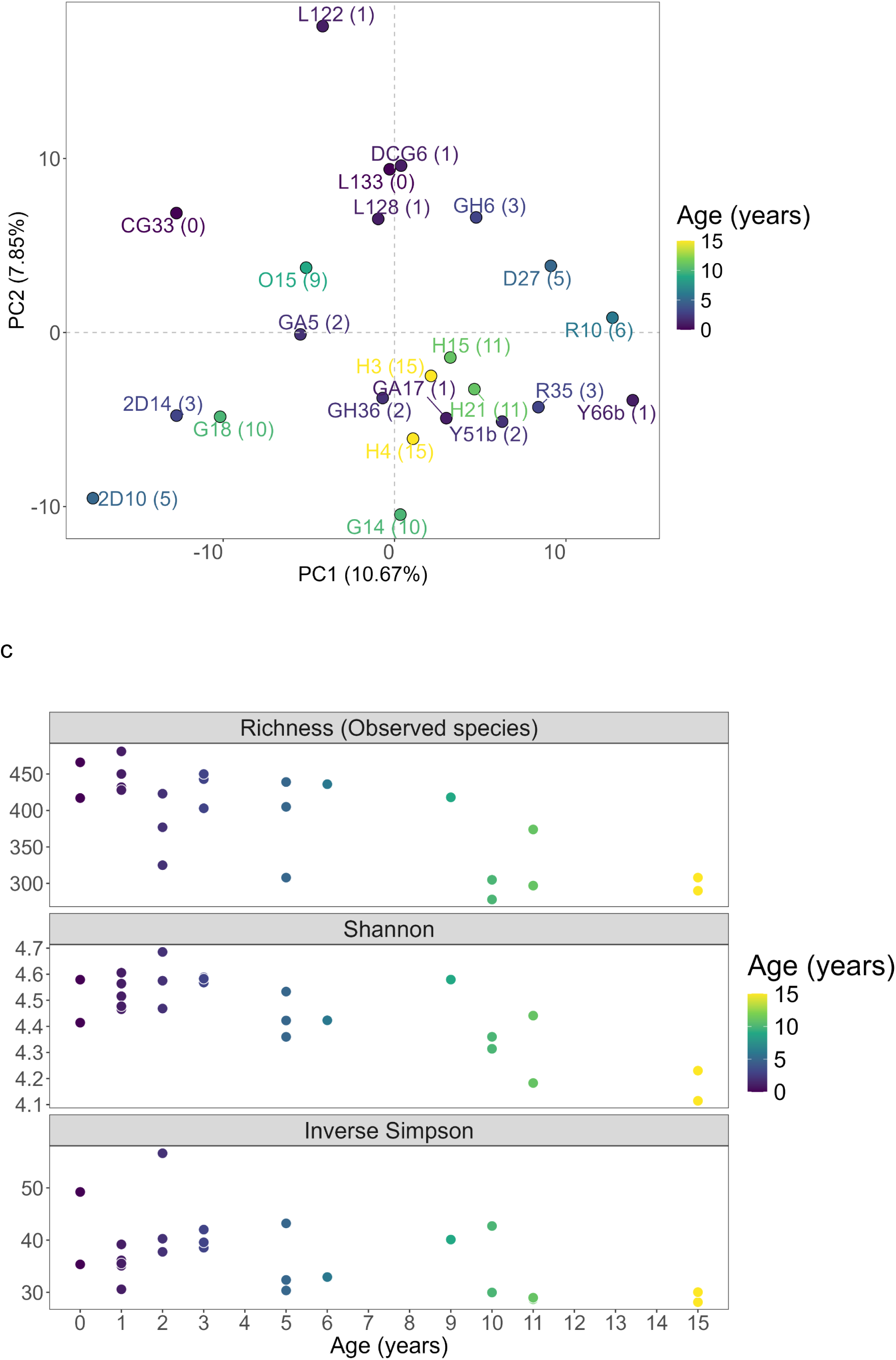
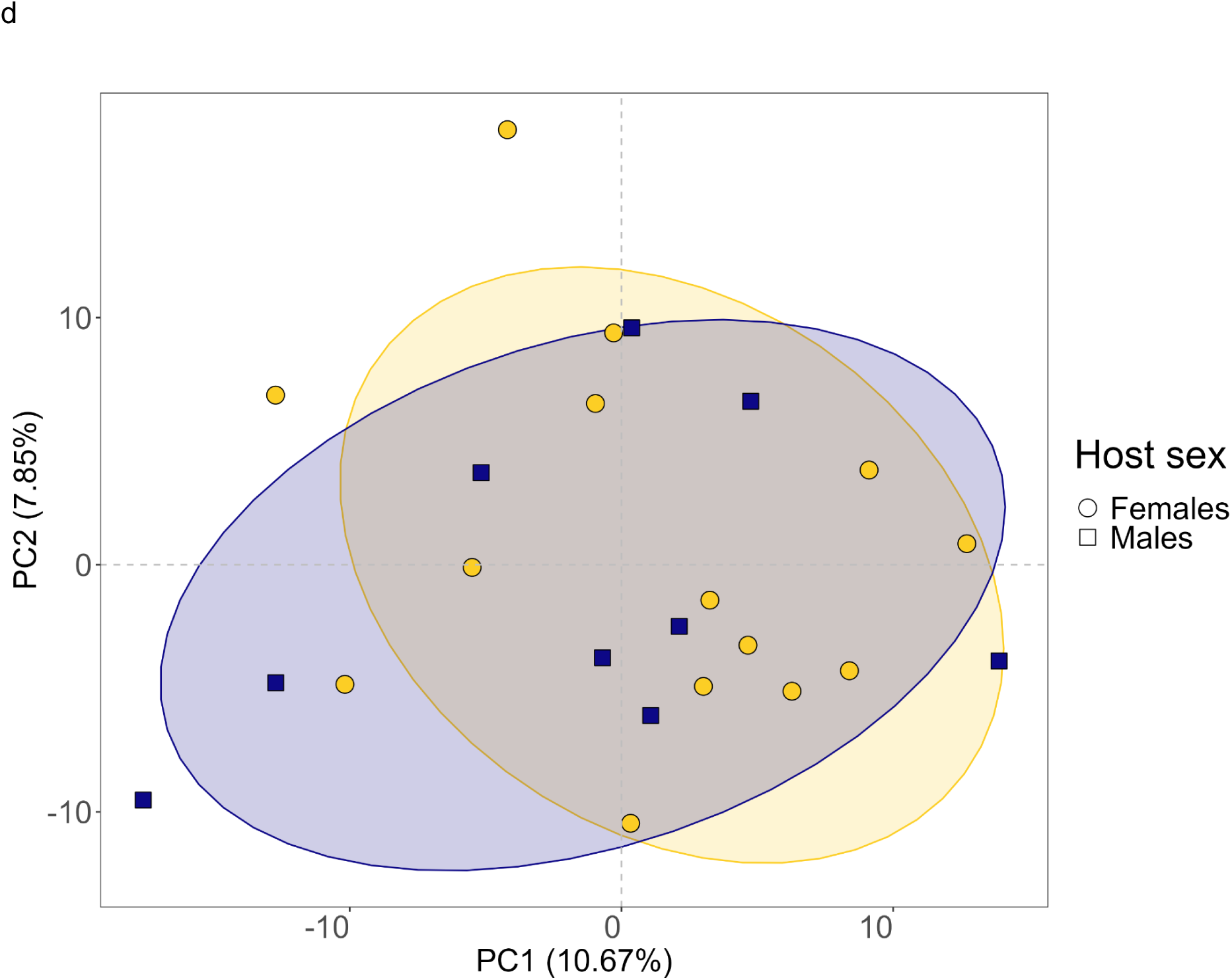
16S rRNA gene sequencing of naked mole-rat gut microbiota, Alpha diversity analysis, and PCA. **a,** The most abundant ASVs across age. Age of each individual is shown in brackets (years). Taxa are abbreviated as follows: Allob.: *Allobaculum*; Erysip. F.: Erysipelotrichaceae Family; Eubac. F.: Eubacteriaceae Family; Fibrob.: *Fibrobacter*; Murib.: Muribaculaceae; Palud. F: Paludibacteraceae Family; Prev.: *Prevotella*; Prevot. F.: Prevotellaceae Family; Prevot. UCG: Prevotellaceae UCG. **b,** There was no separation between old and young naked mole-rat samples. Sample M40 was excluded from PCA on age groups. **c,** Alpha diversity analysis shows that the taxonomic diversity is decreased in old naked mole-rats. **d,** There was no separation between males and females in PCA.

### Whole metagenome sequencing revealed key taxa in the NMR gut

To characterize the gut microbiota of NMRs more closely, whole metagenome sequencing was performed on 11 individuals of different ages (Supplementary Table S12). The Kraken2/Bracken software program could classify 93% of 430M decontaminated reads (average 94% ± 0.4%, n = 11), identifying 34 phyla, 185 families, 343 genera, and 619 species (Figure 4, Supplementary Tables S12, S13, S14, S15, S16).

**Figure 4.**
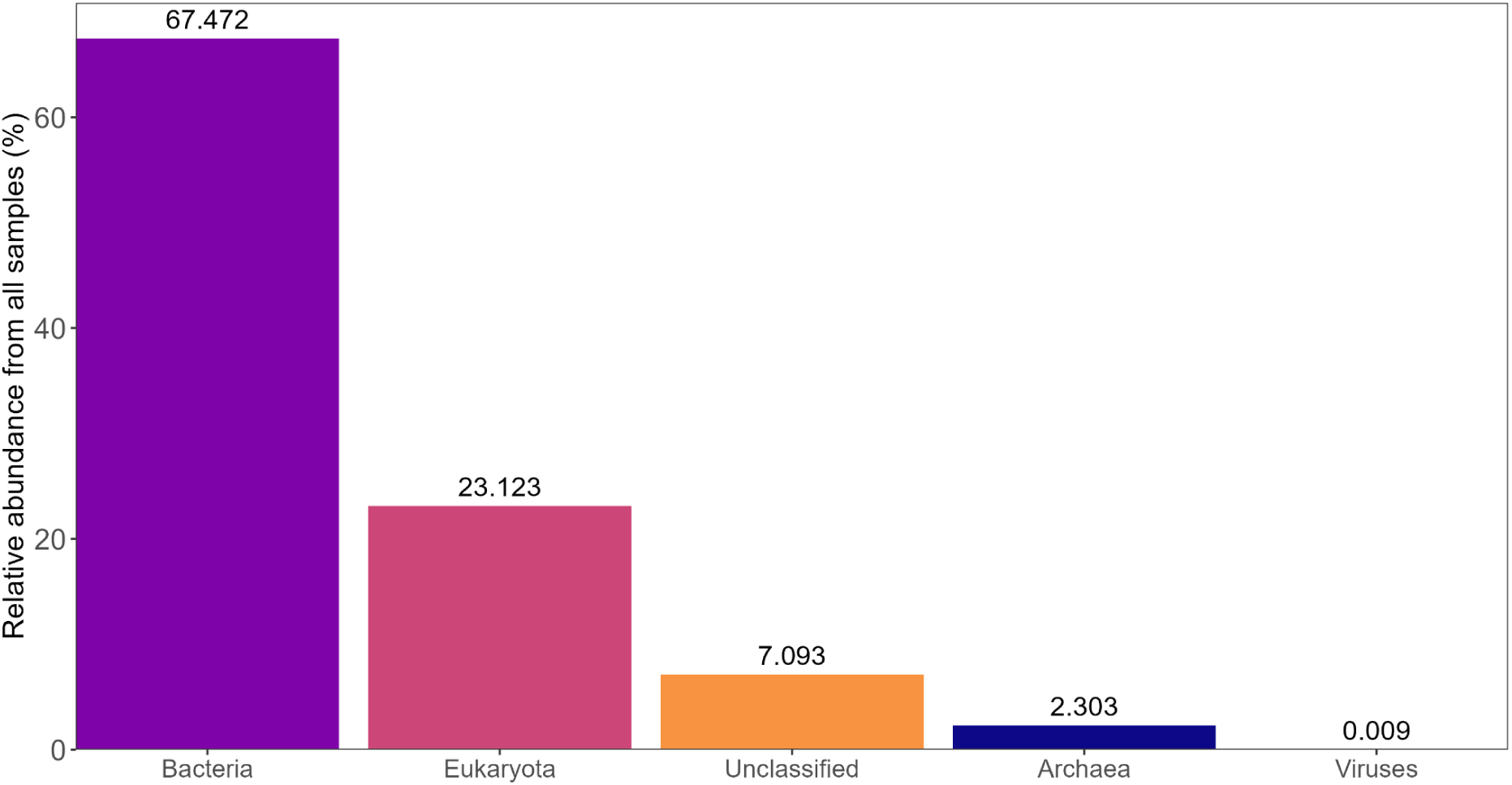
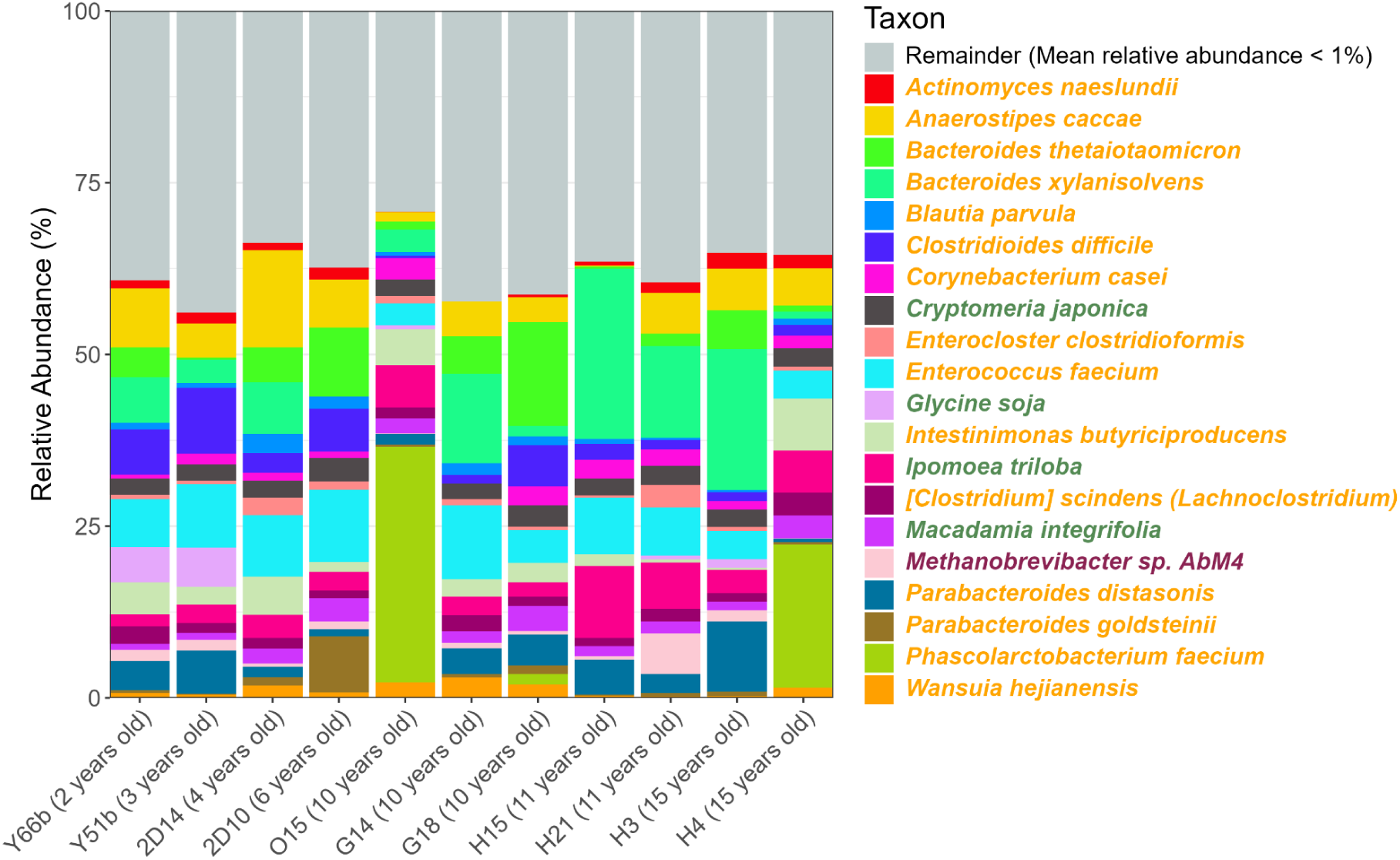
Whole metagenome sequencing profiles. Most reads were attributed to bacteria, dominated by 14 species. Orange names are bacteria, green are eukaryotes, and violet are archaea.

Three bacterial phyla, Bacillota, Bacteroidota, and Actinomycetota, constituted 68.56% of the classified reads (40.5%, 23.2%, and 4.86%) in accordance with the 16S rRNA gene sequencing results (Supplementary Table S17). The most abundant families from Bacillota were Lachnospiraceae (14.4%), Enterococcaceae (7.48%), and Acidaminococcaceae (5.04%); from Bacteroidota were Bacteroidaceae (16.3%), Tannerellaceae (5.31%), and Rikenellaceae (0.778%); and from Actinomycetota were Corynebacteriaceae (2.09%), and Actinomycetaceae (1.7%) (Supplementary Table S18).

*Treponema* has been focused on as the shared taxon between NMR and Hadza, a human rural hunter-gatherer (Debebe et al. 2017), but its average abundance in our whole metagenomic study was 0.021% ± 0.028 (Supplementary Table S19) which is much lower than in our 16S rRNA result (2.2% ± 2.71%). Ten species in *Treponema* were detected, but the most abundant was *Treponema denticola* (0.012%) and the others were negligible (Supplementary figures S4, S5). Moreover, such species were observed only in a few individuals.

We also assessed the abundance of *Desulfobacterota*, which also caught attention in a previous study (Debebe et al. 2017). The abundance of *Desulfovibrio* was lower in the whole metagenome data (0.003%) than in the 16S data (0.385%), and neither *Bilophila* nor *Mailhella*, which can respire using sulfur-containing taurine, was detected in the whole metagenome (Supplementary figures S6, S7). When the gut temperature was predicted using the Metagenomic Thermometer (MetaThermo) based on the amino acid sequences (predicted genes), the result was 28.78 ± 1.59°C, which is best suited for *Desulfovibrio* species.

Within non-bacterial phyla, Streptophyta, Euryarchaeota, and Ascomycota constituted 26.63% of classified reads. The dominant Euryarchaeota family was Methanobacteriaceae (2.48%). The dominant families in Streptophyta were Fabaceae (4.87%), Convolvulaceae (4.49%), Cupressaceae (2.59%), Proteaceae (2.08%), and Salicaceae (1.68%). The dominant families in Ascomycota were Pichiaceae (0.53%), Aspergillaceae (0.36%), and Debaryomycetaceae (0.03%). Apart from Streptophyta, we detected eukaryotic reads from Parabasalia, dominated by Trichomonadidae (0.65%), and Apicomplexa, dominated by Plasmodiidae (0.03%), among others (Supplementary Tables S17, S18).

### MAGs revealed key CAZymes for the tiered fermentation

From 3M contigs with a total length of 4.2 gigabases (Supplementary Table S20), MAGs were created as described below. In the binning process, 1.56M contigs of longer than 2 gigabases were used. After binning, we assembled 1,341 MAGs and filtered them down to 319 high-quality MAGs (completeness ≥90%, contamination ≤5%) (Supplementary Table S21). Dereplication with dRep resulted in 128 representative MAGs (95% ANI) (Supplementary Tables S22, S23). The majority of representative MAGs identified by GTDB-Tk were bacteria (126 MAGs in 11 phyla), and only 2 were archaea (Figure 5a). The most dominant phyla were Bacillota (57 MAGs), Bacteroidota (35), and Spirochaetota (10). The most abundant families were: Lachnospiraceae (17), Acutalibacteraceae (11), and Erysipelotrichaceae (5) in Bacillota; Muribaculaceae (14), Bacteroidaceae (11), and UBA932 (7) in Bacteroidota; and Treponemataceae (8) and Sphaerochaetaceae (2) in Spirochaetota. MAGs in the Bacteroidota and Actinomycetota had relatively large genome sizes, with Bacteroidota having more variable genome lengths (Supplementary figure S8a, Supplementary Table S22). There was a diverse GC% distribution, ranging from 29% in Methanobacteriota to 65% in Desulfobacterota. The most variable GC% were in Bacteroidota (52.8% ± 6.76%) and Bacillota (44.4% ± 9.25%), likely due to the higher number of MAGs in those phyla. We also detected MAGs from Elusimicrobiota, which is characteristic to termites and not rodents. This phylum was also detected in 16S rRNA gene sequencing data (0.23% ± 0.33%) and whole metagenome sequencing data (0.06% ± 0.19%).

**Figure 5.**
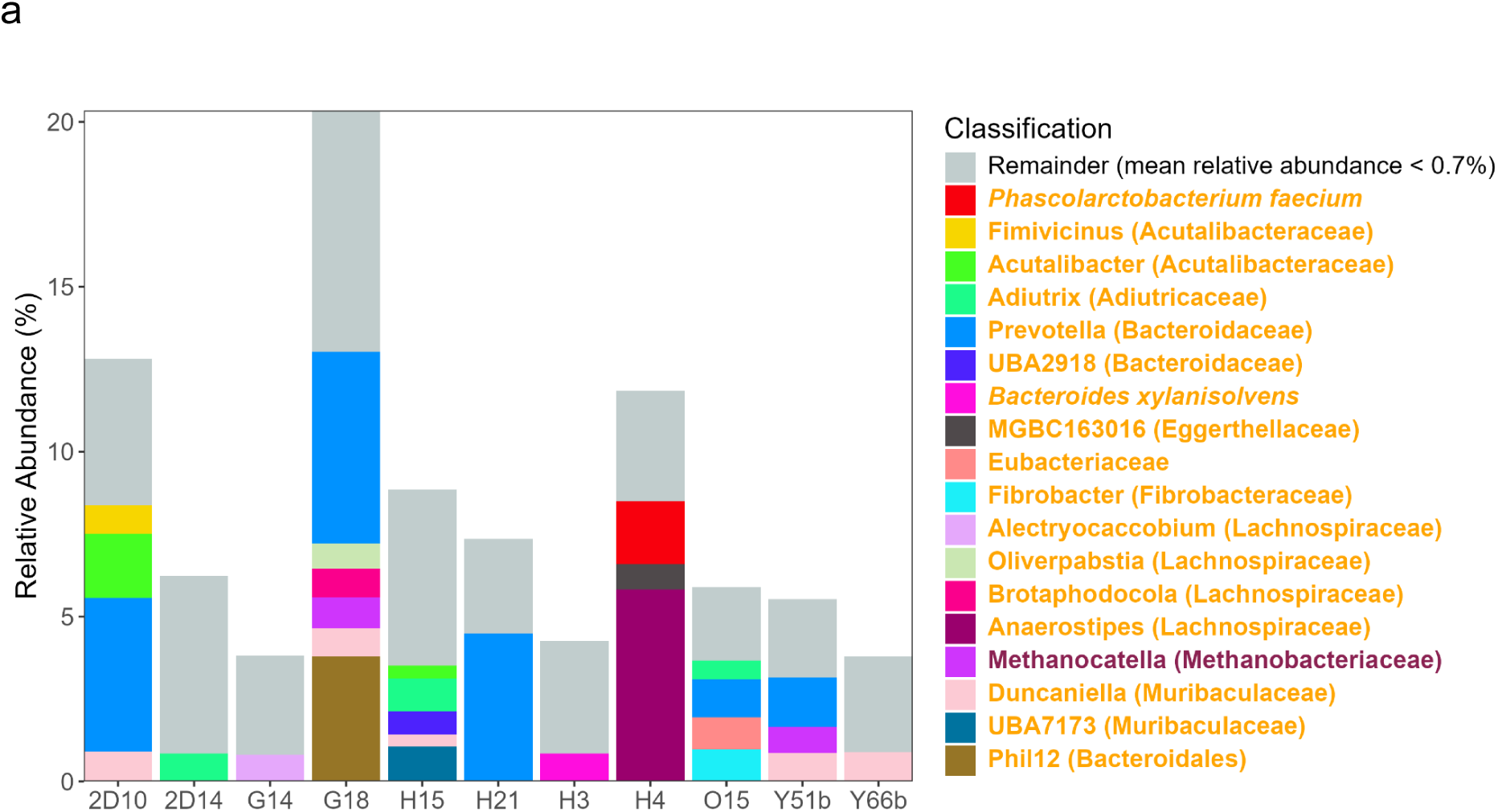
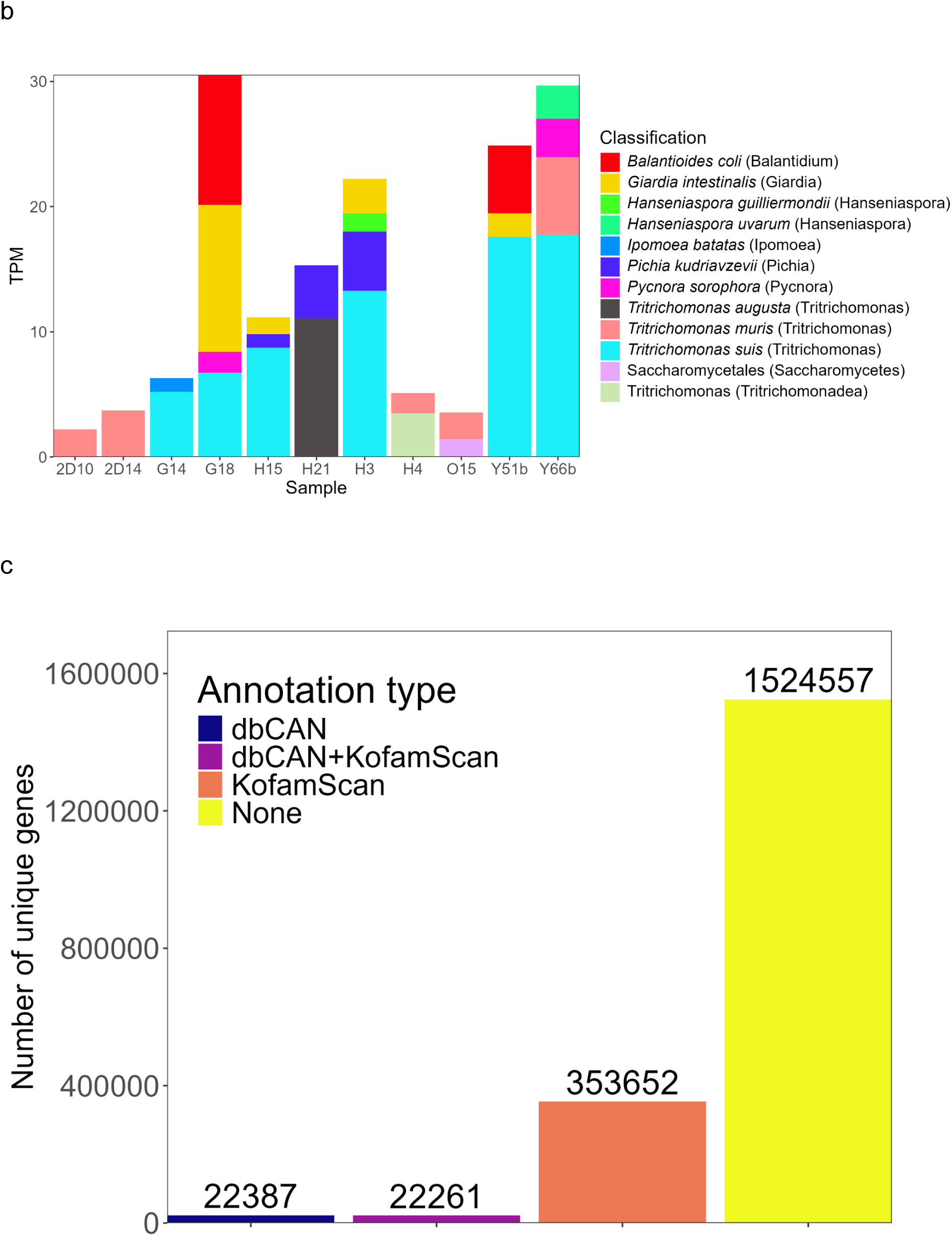
Metagenome assembly and classification results from GTDB-tk, BLASTN, KofamScan, and dbCAN. **a,** MAGs identified by GTDB-tk and **b,** eukaryotic contigs identified by BLASTN (bottom). **c,** Classification of non-redundant proteins by KEGG orthologs (KofamScan) and CAZymes (dbCAN) showed that the majority of coding sequences were unclassified (None).

We also identified 1899 taxa by running BLASTN against the SILVA SSU Ref NR99 database on all contigs (Figure 5b, Supplementary Table S24). 30 of those contigs were eukaryotes, including 8 fungi. Among 8 fungal genera, the dominant ones were *Tritrichomonas* (13 contigs), *Balantidium* (4), and *Giardia* (4). The majority of eukaryotic contigs had a GC% of ∼50%, except *Giardia* (∼75%) and *Balantidium* (30-40%) (Supplementary figure S8c, Supplementary Table S21).

We also built a nonredundant gene catalog containing 1.9M non-redundant coding sequences longer than 100 bp. Gene lengths ranged from 102 bp to 74,937 bp, with a mean length of 582.5 bp and a median length of 405 bp. There were 353,652 proteins assigned to only KEGG Orthology, 22,387 to only Carbohydrate-Active enZymes (CAZymes), 22,261 to both KEGG Orthology and CAZymes, and 1,524,557 not assigned to either database (Figure 5c). After mapping metagenomic reads to genes and quantifying their abundance with Picard, the top 15 genes by median TPM that were present in at least two samples were either unclassified (9 genes) or classified as transposases (6) (Supplementary figure S8d). Among 254 CAZyme subclasses, the top 15 belonged to GH45, GH11, GH44, GH74, GH59, GH98, CBM30, CBM11, CBM88, CBM4, PL14, GT112, and EC 3.2.1.21 (Supplementary figure S8e). CAZymes were mapped to the CAZy database using BLASTP, and mainly belonged to bacteria (40,958) and eukaryotes (3,010). The top hits by bitscore were annotated as *Lactiplantibacillus plantarum* subsp. *plantarum* P-8 (1160 CAZymes), *Carnobacterium maltaromaticum* LMA28 (816), *Arabidopsis arenosa* (696), *Parabacteroides goldsteinii* BFG-241 (632), and *Brassica rapa* (609). GH45 subclass was mapped to *Fibrobacter succinogenes* subsp. *succinogenes* S85 and an uncultured bacterium; GH11 was mapped to an unidentified taxon; GH44 was mapped to *Runella slithyformis* DSM 19594; other top CAZyme subclasses mapped to diverse bacterial taxa and are available in the Supplementary Table S25.

We filtered the list of CAZyme subclasses and KOs to retain core IDs that are shared by more than half of the individuals (at least 5 out of 11 samples). The 256 CAZyme subclasses and 3466 KOs (Supplementary Tables S26, S27) were compared with the core mouse gut CAZyme and KO list from the 2020 Integrated Mouse Gut Metagenome Catalog (iMGMC) (Lesker et al. 2020) to identify IDs not found in the iMGMC data and therefore specific to NMR. The resulting NMR-specific dataset contained 50 CAZyme subclasses and 489 KOs (Supplementary Tables S28, S29).

## Discussion

### Age-associated stability of the NMR gut microbiota

Unlike humans and mice, the microbiota of NMRs remained stable across ages, possibly due to coprophagy and shared environment. No noticeable decrease in the abundance of genera and families was detected (Supplementary Figure S9)(Ghosh et al. 2022), and the differential abundance tests did not reveal any significant differences between sexes or ages (Supplementary Figure S10, S11). This stability may support the unusual longevity of NMRs, but the ages of our samples spanned up to 15 years old, relative to a reported maximum lifespan of 40 years. Continued observations are necessary to confirm the stability of microbiota.

Surprisingly, NMRs seem to maintain their microbial complexity and diversity even after complete rearing in a laboratory environment. In Figure 6, gut microbiota from our laboratory-born individuals are contrasted with those from wild individuals from a separate study by Debebe et al. (2017). The dominant families (Prevotellaceae and Lachnospiraceae) included NMR-specific genera that separate them from the other rodents in our PCA study. Reconstructed MAGs also included many genomes from these families.

**Figure 6.**
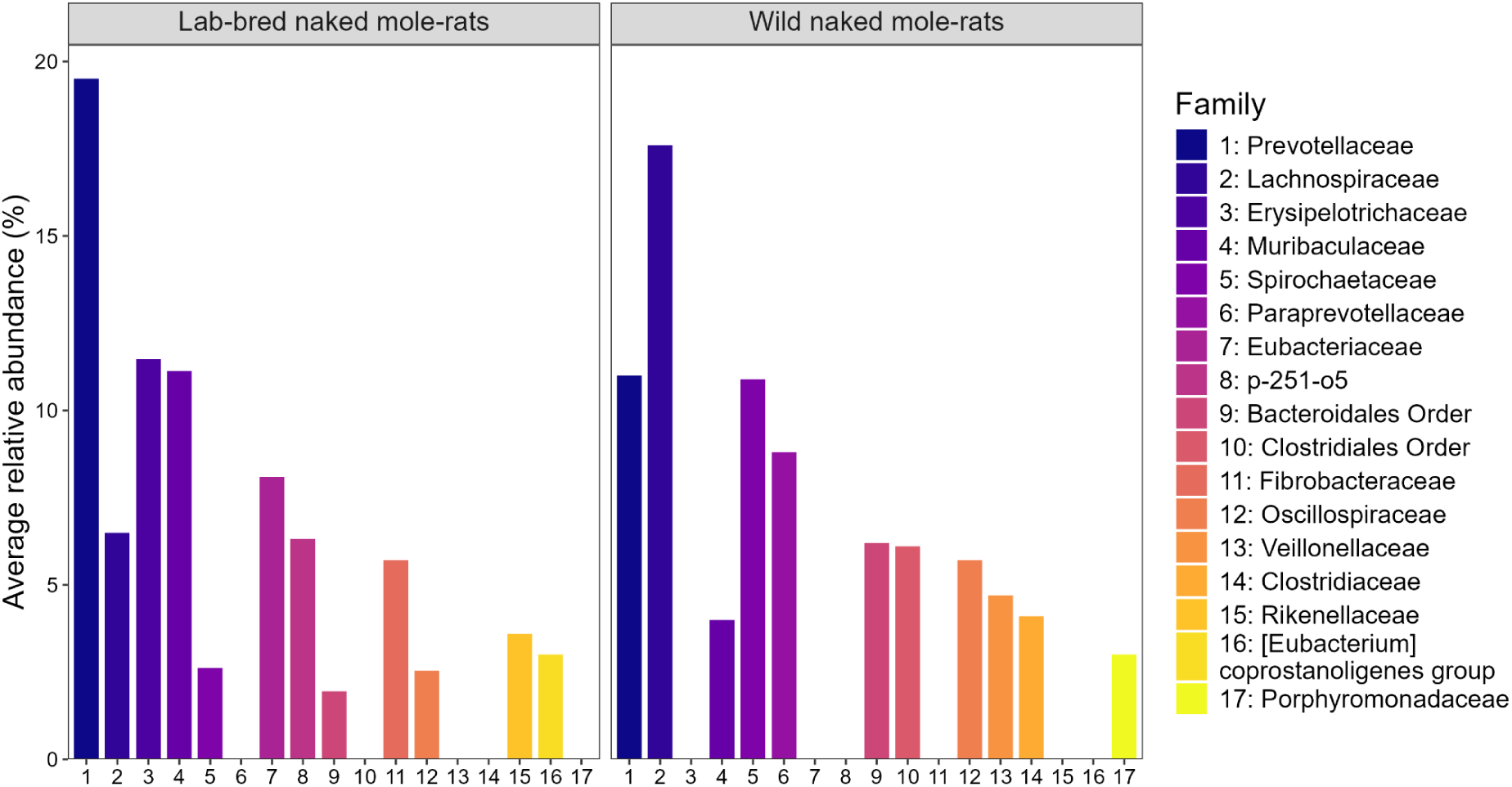
Ten most abundant families in lab-bred and wild naked mole-rats. Data from wild naked mole-rats was taken from the results of the original study (Debebe et al. 2017). Erysipelotrichaceae was mentioned, but no numeric data was provided. The Bacteroidetes S24.7 lineage and Ruminococcaceae were renamed to Muribaculaceae and Oscillospiraceae, respectively. The abundance of Clostridiaceae, Porphyromonadaceae, and Veillonellaceae in laboratory individuals was too low to be shown on the plot (0.00125%, 0.00119%, and 0.000705%, respectively). Paraprevotellaceae was likely a part of Prevotellaceae.

### Tiered degradation of plant polysaccharides

One characteristic genus in the NMR gut is *Treponema*. It has been reported as the dominant genus in the cecum of captive NMRs (Cong et al. 2018), and is also characteristic of wild populations (Debebe et al. 2017). In our metagenomic analysis, the relative abundance of *Treponema* was low; however, 8 out of 128 MAGs were assigned to the family Treponemaceae. Within the 8 genomes, we identified 84 CAZyme subclasses, many of which are involved in plant polysaccharide degradation including glucosidases, xylosidases, and endoglucanases (e.g., GH13, GH3, GH5).

Among the differentially abundant genera identified consistently by two software pipelines was *Fibrobacter*. This genus exists in the foregut of true ruminants, and separated NMR and DMR from other rodents in PCA, accounting for 5.71% ± 5% of the 16S rRNA gene sequencing reads, although it was much less in the whole metagenomic data. One MAG corresponded to *Fibrobacter* and contained 60 CAZyme subclasses. Comparison of CAZyme repertoires between Treponemataceae and *Fibrobacter* MAGs revealed their substantial complementarity: 46 subclasses were unique to Treponemataceae and 22 to *Fibrobacter*. These differences support a model of tiered degradation of plant polysaccharides: *Fibrobacter* serves as a primary degrader characterized by endo-acting CAZymes targeting crystalline cellulose and lignocellulose (e.g., GH45, GH11) together with cellulose-binding modules (CBM30, CBM11) (Suen et al. 2011; Brumm 2013), whereas Treponemaceae species serve as secondary degraders, characterized by CAZymes such as GH5 and GH13 that target partially depolymerised glucans.

Secondary degradation is further supported by two characteristic taxa, p-251-o5 within Bacteroidales and UCG-003 within Prevotellaceae. These taxa were prominent in the 16S rRNA gene sequencing data (6.323% ± 3.728% and 5.481% ± 2.657%, respectively), but were absent in the profiles generated from whole metagenome sequencing due to their absence from the Kraken2 reference database. Nevertheless, BLASTN searches against the SILVA database detected 14 and 32 assembled contigs corresponding to p-251-o5 and UCG-003, respectively. Members of the genus *Prevotella* are well known for their extensive repertoires of CAZymes for polysaccharide degradation.

The members of the family Erysipelotrichaceae, including *Allobaculum*, probably function further downstream in the fermentation. Five MAGs belonged to this family, none of which encoded enzymes for crystalline cellulose; instead, they encoded carbohydrate esterases (CE1) that remove ester substitutions from complex polysaccharides, thereby increasing substrate accessibility, as well as β-glucosidases (GH1) and amylases (GH13) involved in terminal sugar release and secondary processing of soluble or partially solubilized carbohydrates. During fermentation of pectin and hemicellulose by Prevotellaceae and Erysipelotrichaceae, molecular hydrogen can be produced together with short-chain fatty acids such as acetate and propionate.

This hydrogen is likely scavenged by sulfate-reducing bacteria such as *Desulfovibrio*, which were consistently detected in the NMR gut microbiota. Sulfate for this process may be supplied from sulfated mucins in the gut mucus layer as well as from dietary sulfur compounds in plant material. The relatively low abundance of *Desulfovibrio* observed in our data may reflect limited sulfate availability in the diet. Hydrogen can also be consumed by methanogens such as *Methanobrevibacter* and *Methanobacterium*.

In humans, *Methanobrevibacter smithii* is the dominant intestinal methanogen and has been implicated in hydrogen consumption through methanogenesis, thereby influencing bacterial fermentation efficiency and short-chain fatty acid production. Similarly, in mice and true ruminants, *Methanobrevibacter* and *Methanobacterium* species are recurrent members of the gut archaeal community and are thought to modulate microbial metabolic networks by acting as hydrogen sinks.

In contrast to archaeal methanogens, members of the family Paludibacteraceae likely contribute to downstream fermentation through hydrogen-independent pathways, in which reducing equivalents (e.g., NADH) are reoxidized through organic acid formation such as succinate, propionate, and others. In our study, Paludibacteraceae genus was unique to NMRs and was stably detected in the whole metagenomes. Overall, fermentation in the NMR gut appears to rely primarily on sulfate reduction and hydrogen-independent fermentative pathways as hydrogen sinks, likely contributing to substantially lower methane production than that observed in true ruminant mammals.

The naked mole rat–specific core KOs were strongly enriched in functions related to anaerobic and reductive metabolism, including ferredoxin-dependent enzymes, hydrogen metabolism, and archaeal pathways such as methanogenesis. This functional bias suggests that the naked mole rat gut microbiome is adapted to persistently low-oxygen and energy-limited conditions, where efficient redox balancing and hydrogen turnover are critical for maximizing fermentative energy yield. In contrast to mice, whose core gut functions emphasize substrate utilization, the naked mole rat core microbiome appears to be shaped by constraints on electron flow and redox homeostasis, consistent with the host’s subterranean lifestyle and extreme longevity.

The naked mole rat–specific core CAZy repertoire was enriched in carbohydrate-binding modules (CBMs) and glycoside hydrolases (GHs) associated with the deconstruction of insoluble plant cell wall polysaccharides, including cellulose-associated and hemicellulose/pectin-related components. Rather than indicating specialization for cellulose alone, this pattern suggests a coordinated capacity to degrade complex plant cell wall matrices by improving enzymatic access through side-chain removal and polysaccharide modification. This functional configuration is consistent with adaptation to a fiber-rich, energy-limited diet, enabling efficient fermentative energy recovery under the subterranean and hypoxic lifestyle of the naked mole rat.

### Unicellular eukaryotes and thermal signatures

Another notable finding in our study was the detection of unicellular eukaryotic microorganisms that are not captured by 16S rRNA gene sequencing studies. While the Kraken2 pipeline assigned a noticeable fraction of reads to plant species, most likely originating from diets and wooden chip bedding, our metagenome assembly could detect flagellates, ciliates, and fungi. Some of these taxa were tentatively annotated as *Trichomonas*, *Balantidium*, and *Giardia*. The presence of such protozoa is consistent with an early experimental survey (Porter 1957), and members of these taxa are known to play roles in cellulose digestion with their hosts rather than acting solely as parasites. Genus names for protozoa may be different from actual ones because the amount of study is much less compared to standard bacteria and other eukaryotes.

In ruminant mammals, diverse ciliates such as *Diplodinium* actively secrete cellulolytic enzymes and contribute to fiber degradation, while in termites, specialized flagellates such as *Trichonympha* are the primary agents responsible for decomposing lignocellulosic material into fermentable sugars. Protozoa also serve as protein sources for the host; for example, *Isotricha* species functions as rapid scavengers of soluble carbohydrates and bacteria, thereby slowing the fermentation and stabilizing pH of the gut. These well-established systems suggest that protozoa detected in the NMR gut may similarly participate in polysaccharide degradation, potentially contributing to the function of the unusually large cecum in this species.

Interestingly, the predicted gut temperature inferred from gut microbial genes (28.78 ± 1.59°C) closely matched the experimentally measured values by Yahav and Buffenstein (28.9 ± 1.3°C). This prediction was derived solely from the relative frequency of 7 amino acids (I, V, Y, W, R, E, and L) across all encoded proteins, but is known to be surprisingly accurate (Kurokawa et al. 2023). Since the cecum of NMRs has been reported to be slightly cooler than their body temperature, future recovery and analysis of protozoan genomes may allow inference of their preferred microhabitats within the gut based on sequence-derived thermal adaptation signatures.

Members of the phylum Elusimicrobiota have occasionally been detected at low abundance in laboratory mice; however, the genus Avelusimicrobium has rarely, if ever, been reported from murine gut microbiomes. In this study, Avelusimicrobium-affiliated lineages were consistently recovered as high-quality MAGs from naked mole-rat fecal samples across multiple individuals. This pattern suggests that the gut environment of naked mole-rats may support specific Elusimicrobiota lineages that are typically associated with anaerobic, fiber-rich digestive systems, such as those found in insect herbivores. While further comparative and functional analyses will be required to determine the extent of host specificity, the recurrent presence of Avelusimicrobium indicates that it may represent a distinctive component of a gut microbiome shaped by the unique ecology and diet of naked mole-rats.

Taken together, we performed the first whole metagenome sequencing of the NMR gut microbiota, providing sequence-based evidence for microbial diversity and revealing its metabolic potential. Our high-quality dataset now enables linking novel metabolic genes with host physiology and ecological niche. For example, assessing the actual enzyme activities of Treponema may reveal similar results to nitrogen fixation in termites (Lilburn et al. 2001; Graber et al. 2004), although it requires isolation and cultivation of bacteria as described by Evans et al. (Evans et al. 2011). The ultimate challenge is establishing a causal relationship between the metabolism of novel taxa and the stability of the NMR’s microbiota, which will help us understand how the gut microbiota contributes to its ‘healthy’ longevity.

## Acknowledgments

We are grateful to Prof. Kirill Kryukov for advice on metagenome analysis, and to Dr. Masanori Yamakawa for fruitful discussions on naked mole-rat behaviour. We thank M. Kobe and M. Nagata for animal maintenance; Y. Tanabe for laboratory maintenance; and all members of M.A. and K.M. laboratories for technical assistance and scientific discussion. This work was supported in part by MEXT KAKENHI (MA 23H02470; HB 22H03149; KM 21H02392, 22K18355, 23K20043, 24H00542 and 25K21744; KO 22K15024, JP25K09477; YK 22K06069, 23H04948, 25K02197), JST FOREST (KM JPMJFR216C), JST COI-NEXT (KM JPMJPF2010), Naito Foundation to KM and KO, Soroptimist Foundation Japan to KM, Takeda Science Foundation to KM, Toray Science Foundation to YK, and KOSE Cosmetology Research Foundation to KO, and the SOKENDAI Internship Program (2023). Computations were performed on the NIG Supercomputer at the ROIS National Institute of Genetics.

## Data availability

The raw 16S rRNA gene sequences and whole metagenome sequences are available at PRJNA1405902. Processed data is available at Zenodo (https://doi.org/10.5281/zenodo.18454971).

## Notes

https://doi.org/10.5281/zenodo.18454971

https://www.ncbi.nlm.nih.gov/bioproject/PRJNA1405902

### Competing Interest Statement

The authors have declared no competing interest.

