## Supplemental figures for "Diversity and stability of the gut microbiome of naked mole-rat (*Heterocephalus glaber*), the longest-lived rodent"

### Supplementary methods, results, figures, and references

#### List of figures

Supplementary figure S1: Abundance of bacterial genera in the 16S rRNA gene sequencing data (mean relative abundance ≥ 1% within a host).

Supplementary figure S2: Differentially abundant bacteria that were characteristic of naked mole-rats (16S rRNA gene sequencing data)

Supplementary figure S3: Alpha diversity analysis showed no differences between male and female naked mole-rats (16S rRNA gene sequencing data).

Supplementary figure S4: Abundance of the *Treponema* genus in the whole metagenome sequencing data.

Supplementary figure S5: Abundance of genera from the Spirochaetota phylum (including *Treponema*) in the 16S rRNA gene sequencing data.

Supplementary figure S6: Abundance of the Desulfobacterota phylum in the whole metagenome sequencing data.

Supplementary figure S7: Abundance of genera from the Desulfobacterota phylum in the 16S rRNA gene sequencing data.

Supplementary figure S8: Metagenome assembly and classification results from GTDB-tk, BLASTN, KofamScan, and dbCAN.

Supplementary figure S9: Relative abundance of Group 1, 2, and 3 members described in Ghosh et al (2022) from whole metagenome sequencing.

Supplementary figure S10: Only three ASVs were differentially abundant in young individuals (16S rRNA gene sequencing data).

Supplementary figure S11. *Mageeibacillus indolicus* was significantly more abundant in young naked mole-rats in the whole metagenome sequencing data.

### Supplementary methods

#### Differential microbiome analysis

MaAslin2 used a table of absolute abundances with min_prevalence=0, the LM analysis method, TSS normalization, the LOG transform, and a significance threshold of 0.05.

For ALDEx2 analysis, a model matrix was provided in addition to the absolute abundance table. The model matrix contains samples as rows and tested metadata variables (animal hosts) used for the differential abundance test as columns. A value in the matrix is “1” if a sample is from the reference host and “0” otherwise. The aldex.clr function used 1000 Monte-Carlo instances, while all other parameters were set to default. Significant features were those with an absolute value >1 and whose confidence interval did not overlap with zero.

ANCOM-BC used a phyloseq object with the absolute abundance table in a wide format, a formula that specifies tested metadata variables (animal hosts), a method to adjust p-values (FDR), a prevalence filter set to 0 (don’t exclude rare taxa), library size (number of reads in a sample) threshold (set to 0), group parameter set to class, struc_zero=TRUE, neg_lb=TRUE, tol=1e-5, conserve=TRUE, max_iter=100, alpha=0.05, and global=TRUE.

### Supplementary results

#### 16S rRNA gene sequencing analysis

As for DMR data, the original report analyzed the sequences from frozen and freeze-dried samples. However, we used only frozen samples in our re-analysis. Therefore, the numbers in our subset became less than the original, whole dataset: our subset samples contained 19 phyla, 97 (instead of 117) families, and 165 (instead of 210) genera (124 classified, 41 unclassified), and 1,368 ASVs (instead of 1,768).

The abundance of dominating phyla was consistent with the original analysis: The mean relative abundances for Bacteroidota and Bacillota were 69.7% (range 41.8-90.4%) and 22.5% (range 4.01-40.4%) in our analysis, and 68.9% (31.4-86.7) and 20.3% (5.2-47.8) in the original.

The authors of the PVO paper clustered ASVs to 97%-identity OTUs, which makes comparison with our re-analysis difficult. Apart from the nine major phyla, the authors did not provide the number of classified families and genera. However, they mentioned 12 dominant families in the core microbiota. The authors obtained 1,104 OTUs, of which 683 OTUs belonged to PVO. They generated 3,665,256 high-quality paired-end sequences, with an average library size of 9,561 non-chimeric reads/sample (range 959 – 33,265 reads/sample). However, they did not provide the read number and library size for PVO specifically. We did not find the original ASV table from the paper, either. The total read number in the paper is higher than in our analysis (758,432).

The major phyla in the flying squirrel samples were Firmicutes (62.4-100%) and Bacteroidota (0-19.29%), which were slightly different from our data. Firmicutes range was 27.8-75.4%, whereas Bacteroidota range was 15.0-31.4%. The 12 dominant families in the original paper included Lachnospiraceae (50.87%), Ruminococcaceae (28.475%), and Muribaculaceae (8.287%). These three families were also major in our analysis: Lachnospiraceae (mean 36.6%), Muribaculaceae (15.3%), Ruminococcaceae (9.2%). The results differ partly because we excluded nine samples due to low abundance. Nevertheless, they are consistent with each other.

The authors of the hare and rabbit paper identified 14 bacterial phyla, 83 families, 70 genera (58 classified, 12 unclassified), and 6,662 ASVs (Supplementary data of the original paper). The total abundance was 14,559,822 reads (mean 501,908 reads per sample, excluding control samples). Although we removed pregnant and lactating hosts from re-analysis, we identified more families (120) and genera (241), but fewer ASVs (4642).

The dominant phyla coincided in both the original and our results. Firmicutes and Bacteroidetes together comprised >90% of the community in our results. Consistent with the original data, Firmicutes were mainly represented by Clostridiaceae, Lachnospiraceae, Ruminococcaceae, and Oscillospiraceae, while Bacteroidetes were represented by Bacteroidaceae, unclassified Bacteroidales, Rikenellaceae, and Muribaculaceae.

#### Differential abundance tests within naked mole-rats

The differential abundance test with MaAsLin2 confirmed the stability of NMR microbiota. Only three ASVs (*Lactobacillus*, *Ligilactobacillus,* and *Streptococcus*) were characteristic of young naked mole-rats, and no ASVs were detected as significant when comparing males and females.

When comparing the gut microbiota composition of young and old naked mole-rats in the Kraken2 data, MaAsLin2 detected only one differentially abundant species. The characteristic bacterium for young naked mole-rats was *Mageeibacillus indolicus* (p = 0.003, average 0.008% for all individuals, 0.0219% for the young group, 0.0009% for the old group)(Figure S11).

### Supplementary figures

#### Figure S1:

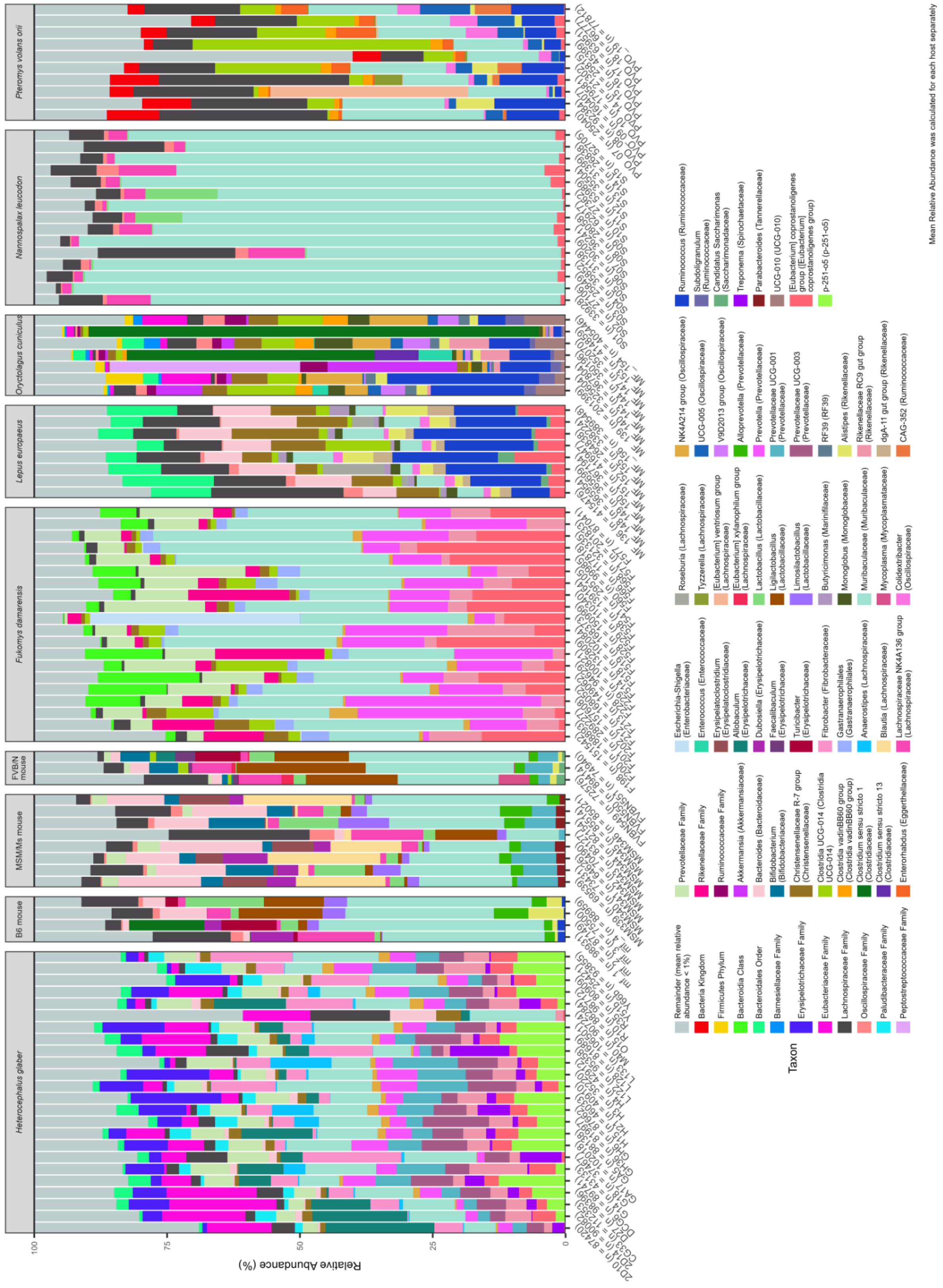

**Figure S1 | Abundance of bacterial genera in the 16S rRNA gene sequencing data (mean relative abundance ≥ 1% within a host).**

Figure S2:

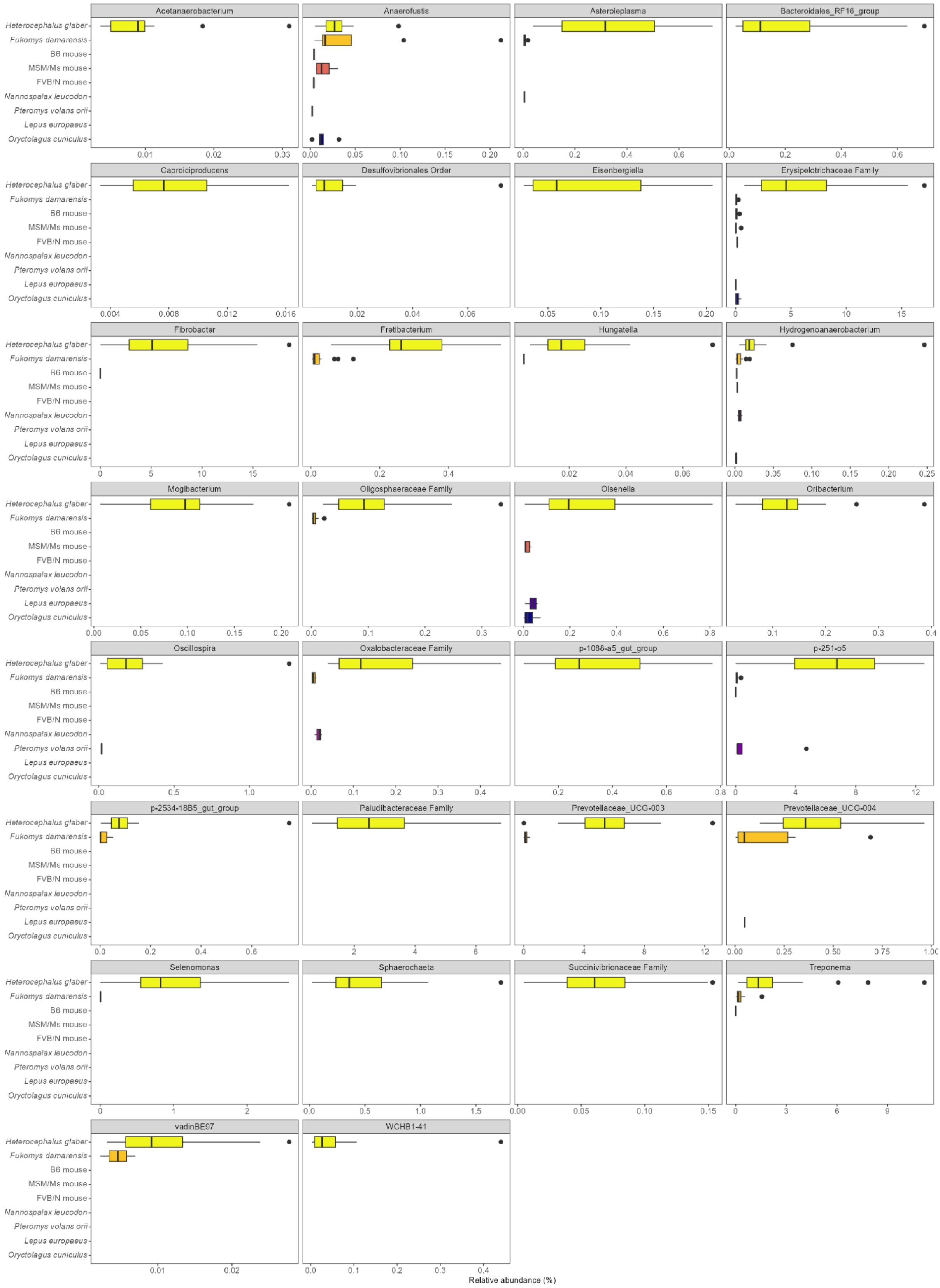
Acetanaerobacterium
Anaerofustis
Asteroleplasma
Bacteroidales_RF16_group
Caproiciproducens
Desulfovibrionales Order
Eisenbergiella
Erysipelotrichaceae Family
Fibrobacter
Fretibacterium
Hungatella
Hydrogenoanaerobacterium
Mogibacterium
Oligosphaeraceae Family
Olsenella
Oribacterium
Oscillospira
Oxalobacteraceae Family
p-1088-a5_gut_group
p-251-o5
p-2534-18B5_gut_group
Paludibacteraceae Family
Prevotellaceae_UCG-003
Prevotellaceae_UCG-004
Selenomonas
Sphaerochaeta
Succinivibrionaceae Family
Treponema
vadinBE97
WCHB1-41
Heterocephalus glaber
Fukomys damarensis
B6 mouse
MSM/Ms mouse
FVB/N mouse
Nannospalax leucodon
Pteromys volans orii
Lepus europaeus
Oryctolagus cuniculus
Relative abundance (%)

**Figure S2 | Differentially abundant bacteria that were characteristic of naked mole-rats (16S rRNA gene sequencing data).**

Figure S3:

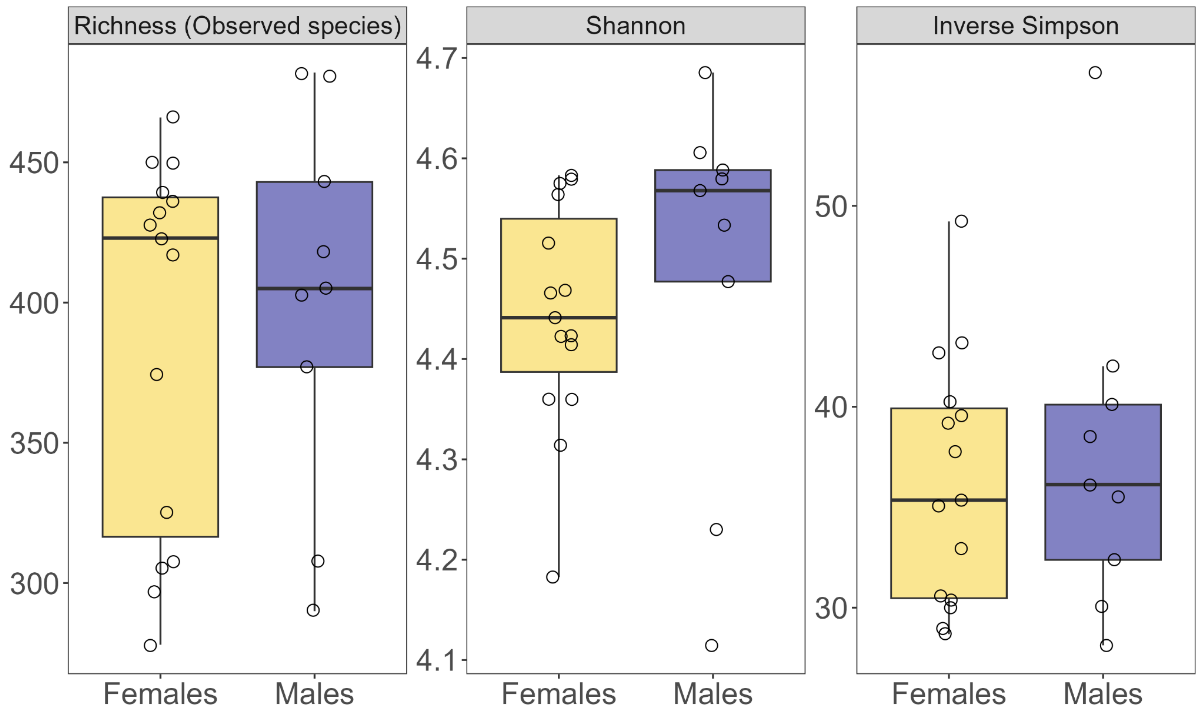


**Figure S3 | Alpha diversity analysis showed no differences between male and female naked mole-rats (16S rRNA gene sequencing data).**

Figure S4:

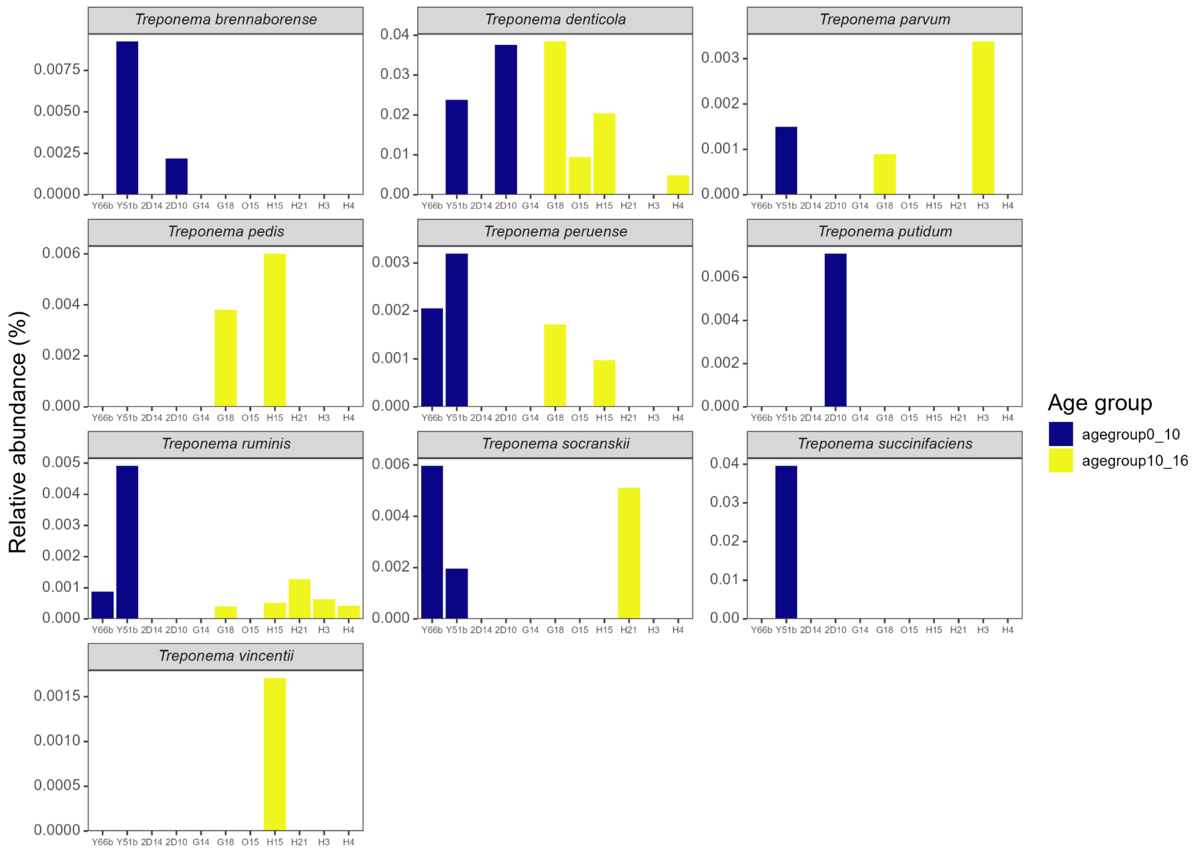


**Figure S4 | Abundance of the *Treponema* genus in the whole metagenome sequencing data.**

Figure S5

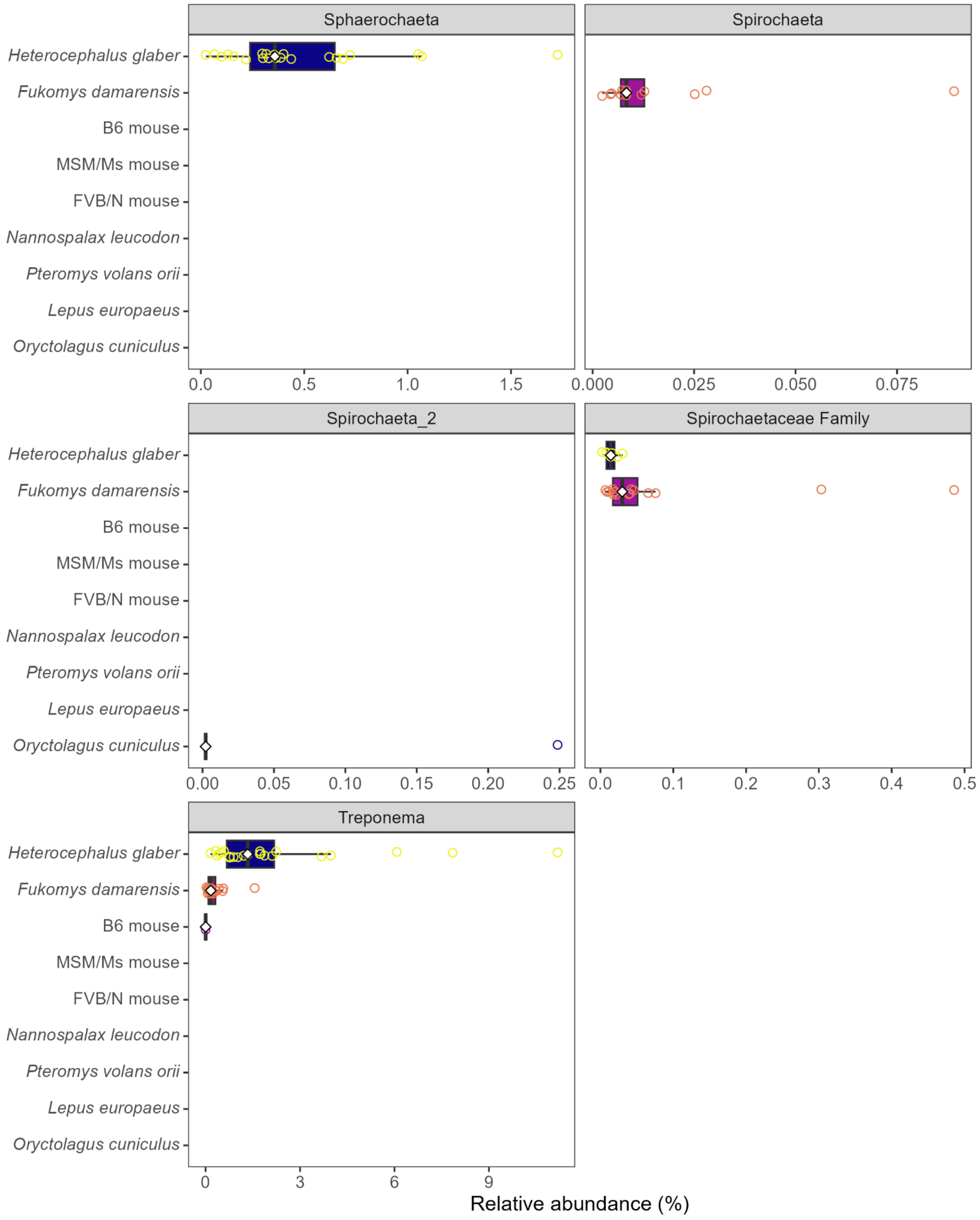


**Figure S5 | Abundance of genera from the Spirochaetota phylum (including *Treponema*) in the 16S rRNA gene sequencing data.**

Figure S6:

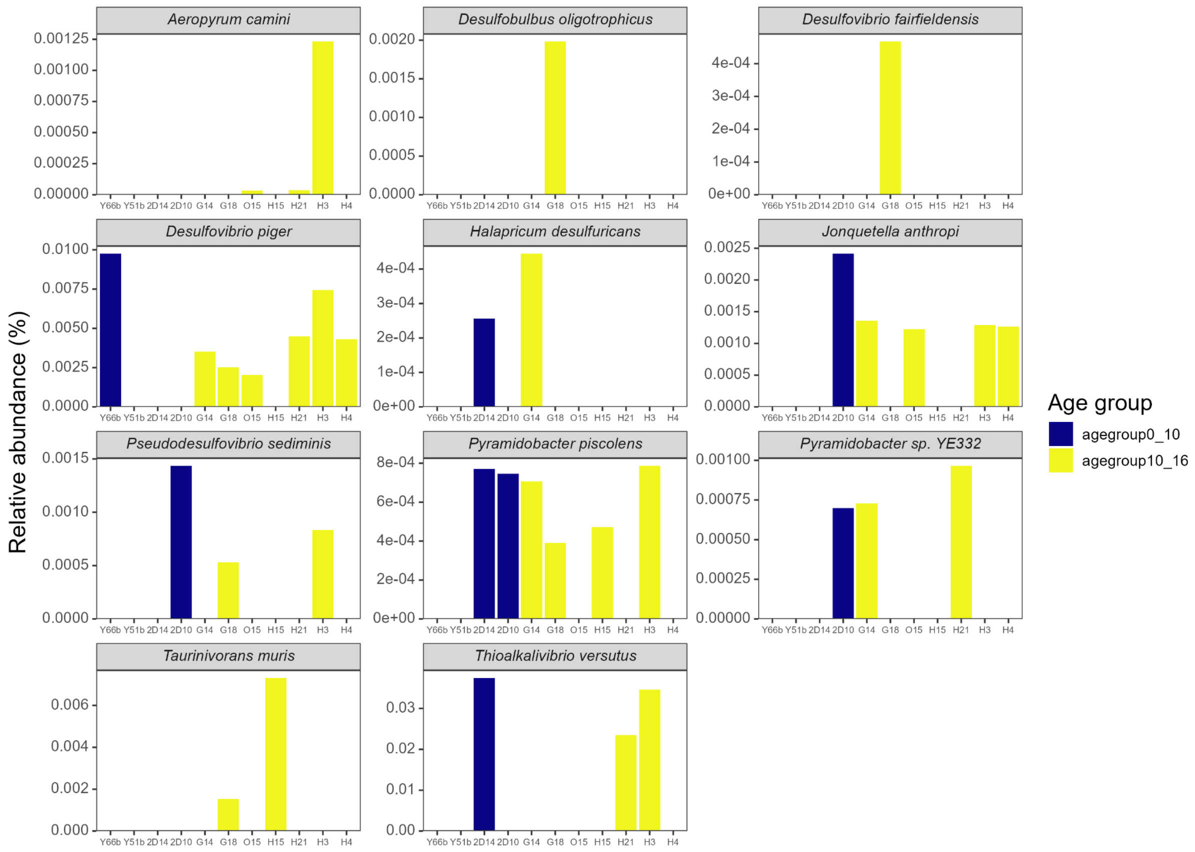


**Figure S6 | Abundance of the Desulfobacterota phylum in the whole metagenome sequencing data.**

Figure S7:

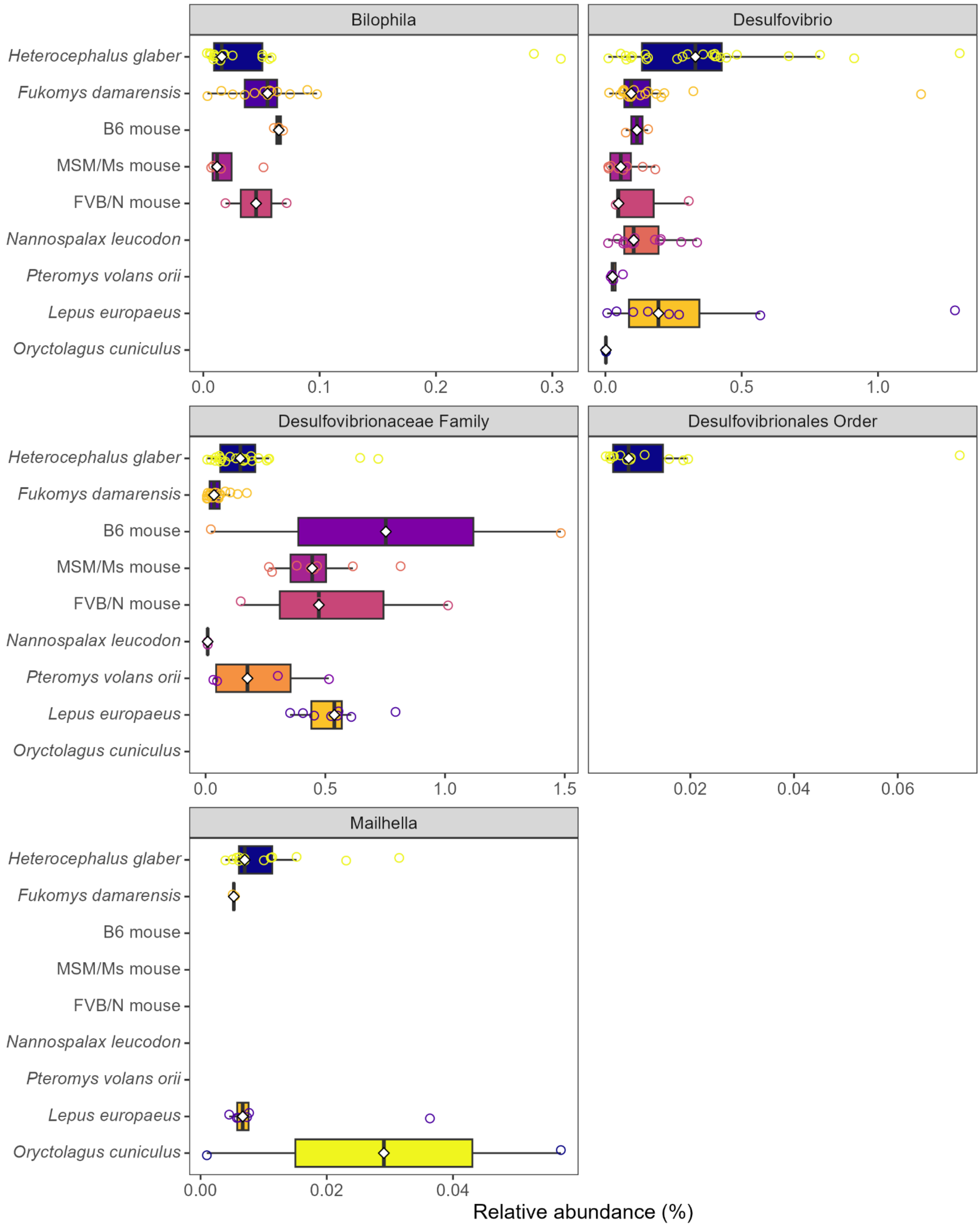


**Figure S7 | Abundance of genera from the Desulfobacterota phylum in the 16S rRNA gene sequencing data.**

Figure S8:

a

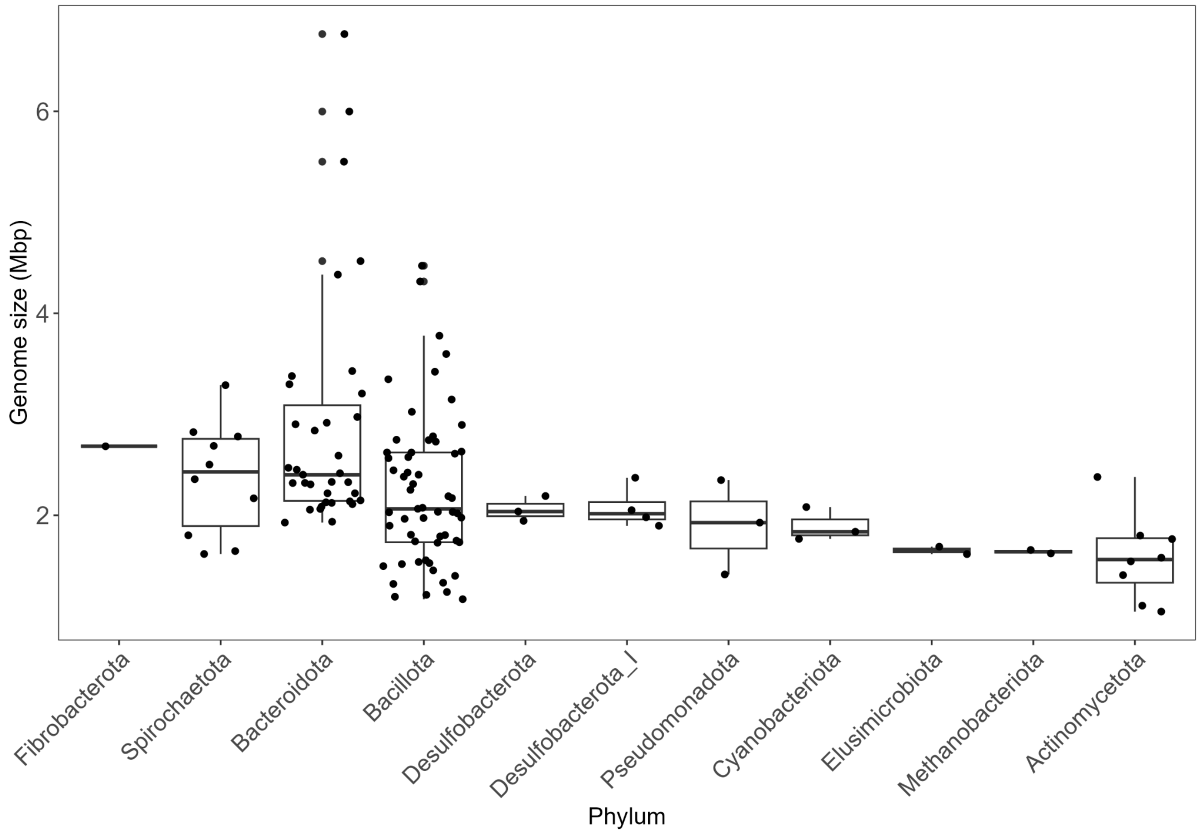

Genome size (Mbp)
6
4
2
Fibrobacterota
Spirochaetota
Bacteroidota
Bacillota
Desulfobacterota
Desulfobacterota_I
Pseudomonadota
Cyanobacteriota
Elusimicrobiota
Methanobacteriota
Actinomycetota
Phylum


b

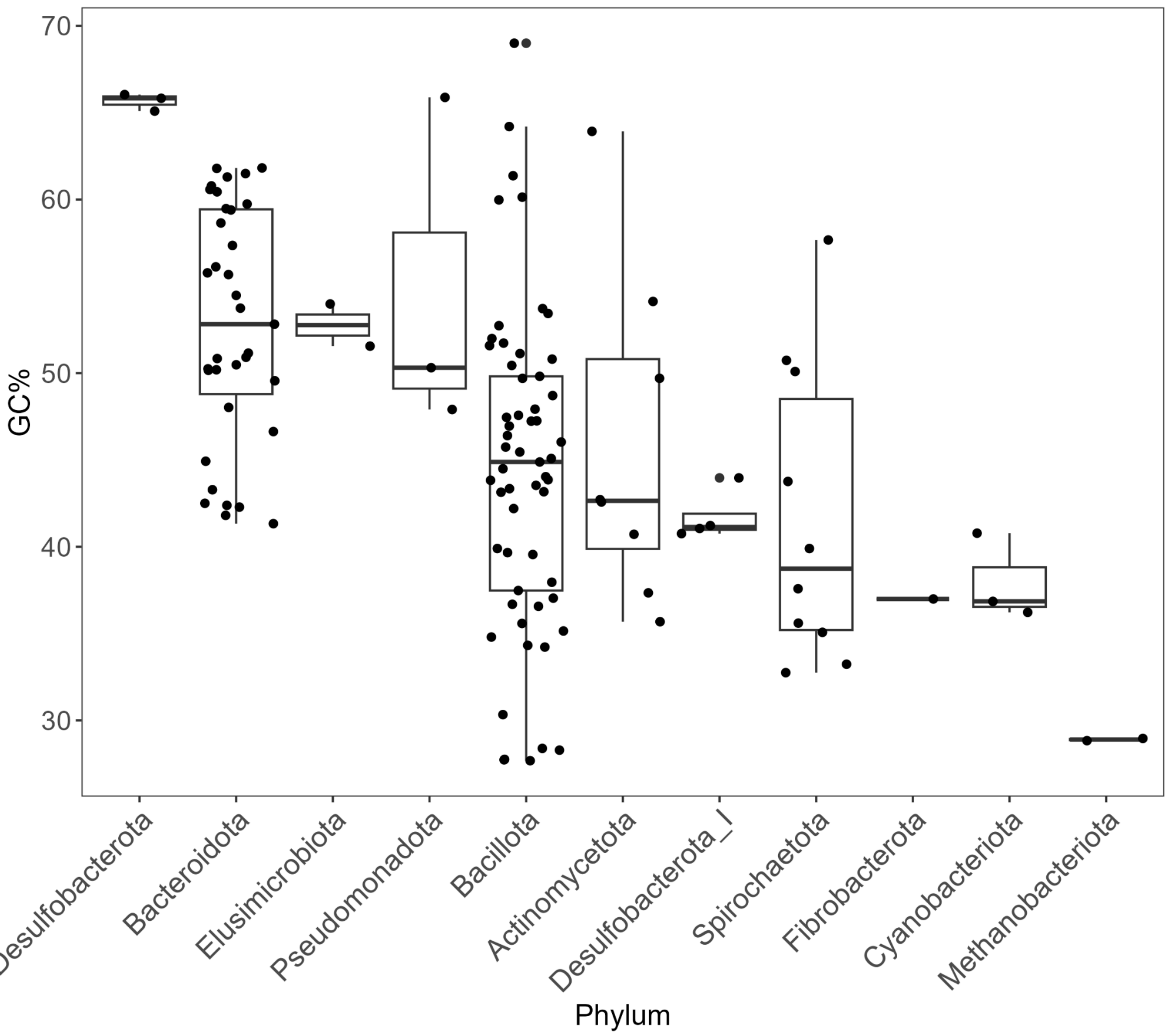

70
60
50
40
30
GC%
Desulfobacterota
Bacteroidota
Elusimicrobiota
Pseudomonadota
Bacillota
Actinomycetota
Desulfobacterota_I
Spirochaetota
Fibrobacterota
Cyanobacteriota
Methanobacteriota
Phylum


c

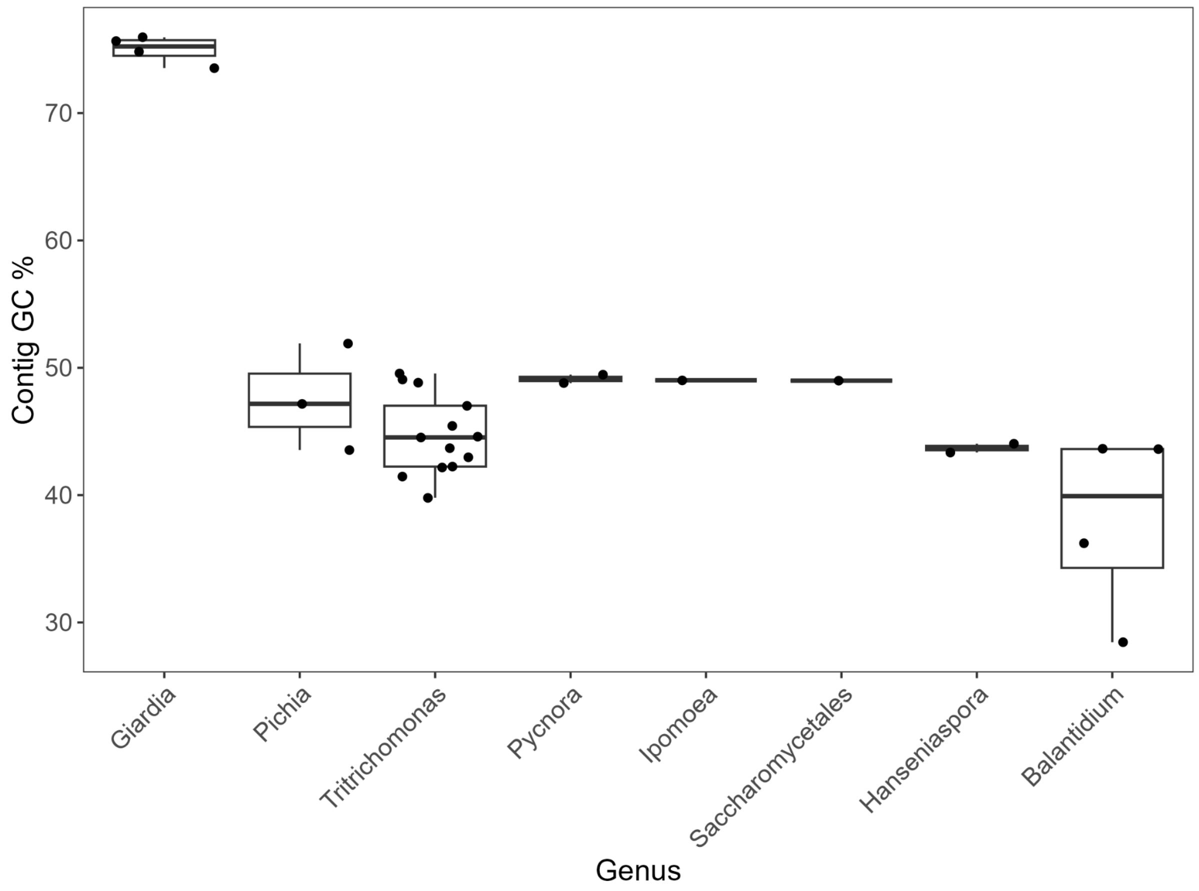
Contig GC %
70
60
50
40
30
Giardia
Pichia
Tritrichomonas
Pycnora
Ipomoea
Saccharomycetales
Hanseniaspora
Balantidium
Genus

d

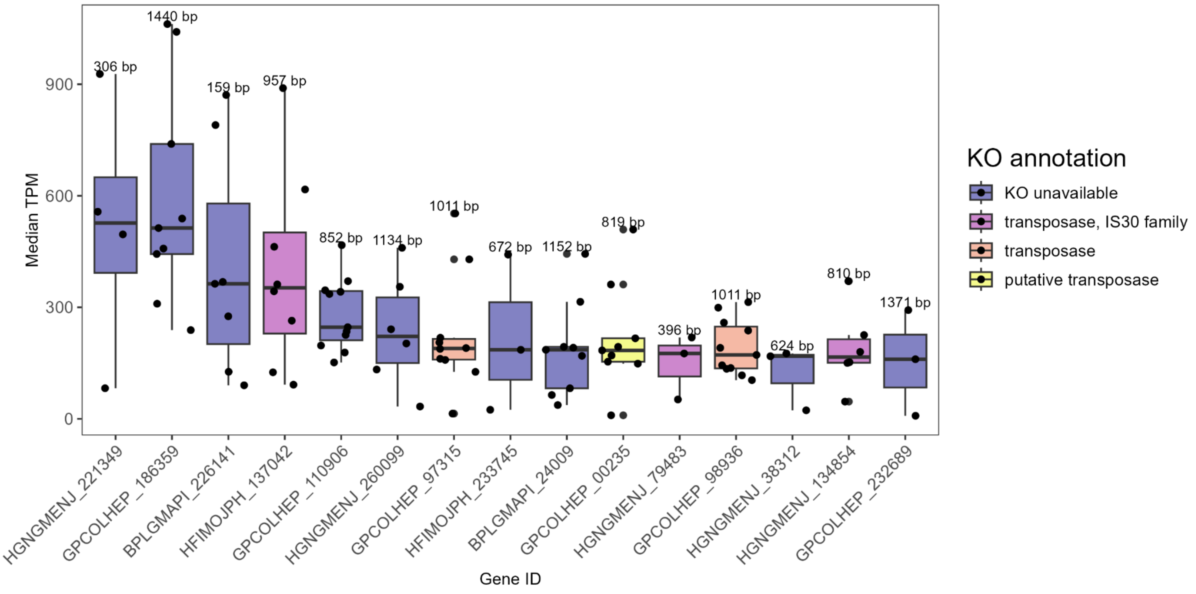
Median TPM
900
600
300
0
306 bp
1440 bp
159 bp
957 bp
852 bp
1134 bp
1011 bp
672 bp
1152 bp
819 bp
396 bp
1011 bp
624 bp
810 bp
1371 bp
HGNGMENJ_221349
GPCOLHEP_186359
BPLGMAPI_226141
HFIMOJPH_137042
GPCOLHEP_110906
HGNGMENJ_260099
GPCOLHEP_97315
HFIMOJPH_233745
BPLGMAPI_24009
GPCOLHEP_00235
HGNGMENJ_79483
GPCOLHEP_98936
HGNGMENJ_38312
HGNGMENJ_134854
GPCOLHEP_232689
Gene ID
KO annotation
KO unavailable
transposase, IS30 family
transposase
putative transposase

e

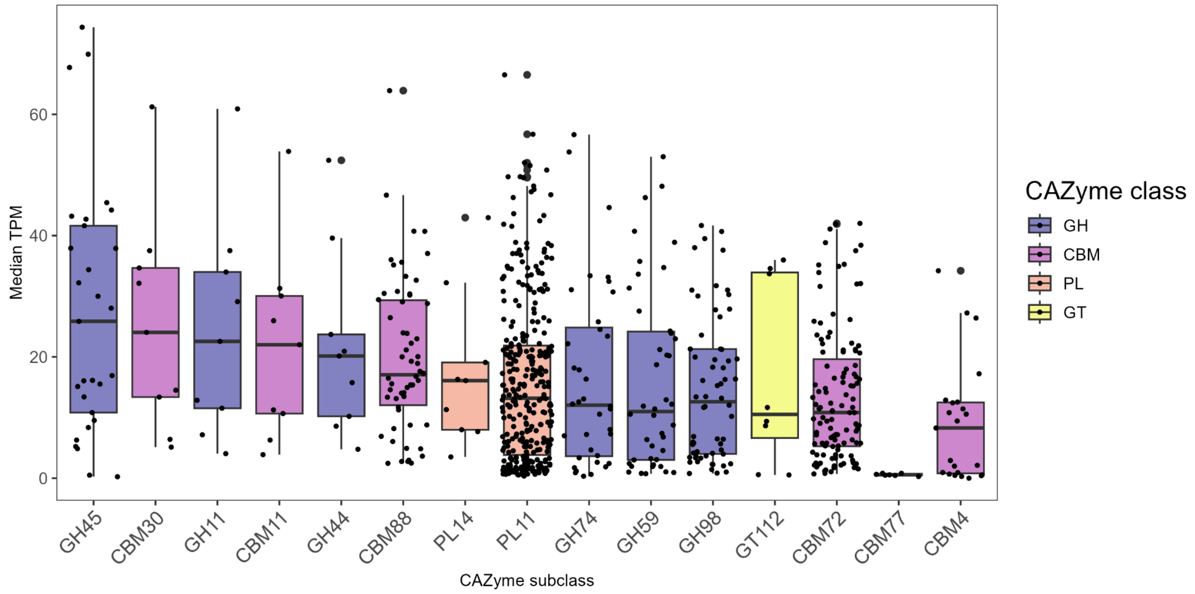


**Figure S8 | Metagenome assembly and classification results from GTDB-tk, BLASTN, KofamScan, and dbCAN. a,** MAG genome lengths. **b,** GC% distribution in MAG phyla and **c,** eukaryotic genera. **d,** Top 15 genes by TPM found in at least 2 samples. **e,** top 15 CAZyme subclasses in all data, including unassembled contigs.

#### Figure S9:

a

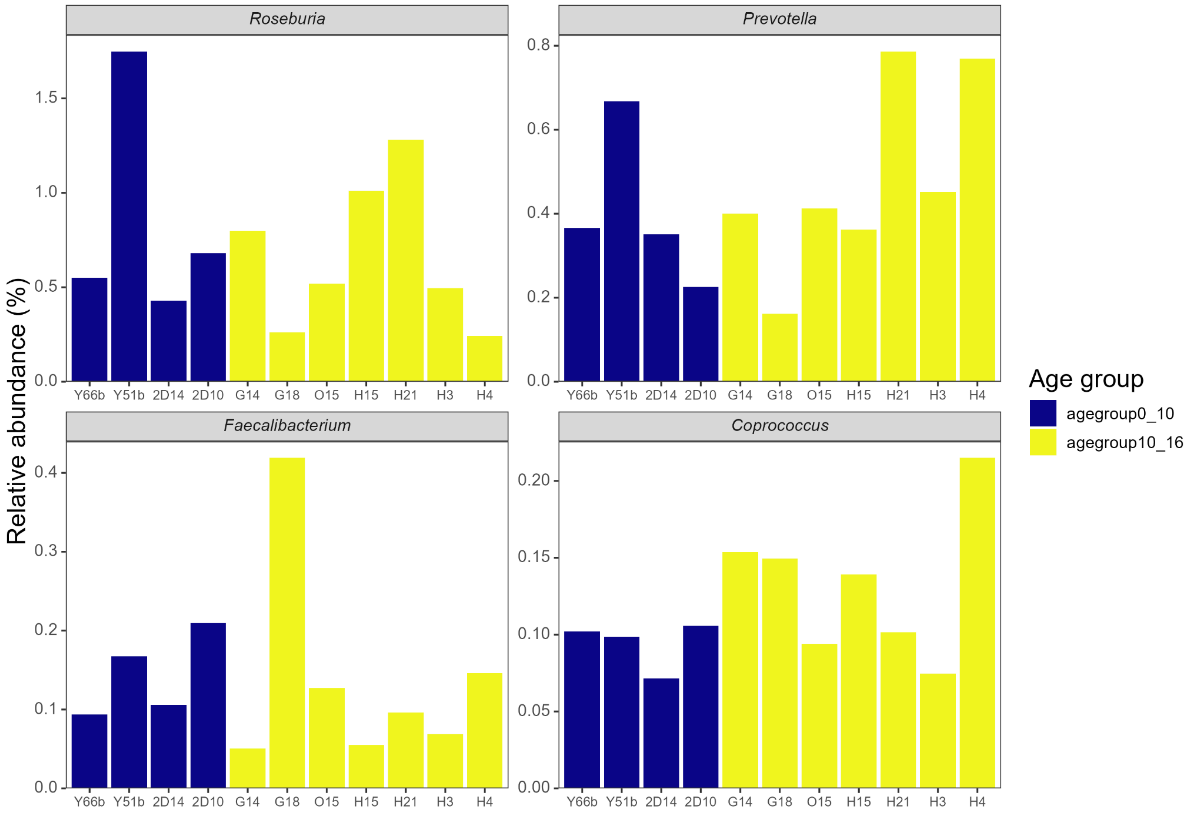
Roseburia
Prevotella
Faecalibacterium
Coprococcus
Relative abundance (%)
Y66b Y51b 2D14 2D10 G14 G18 O15 H15 H21 H3 H4
Age group
agegroup0_10
agegroup10_16

b

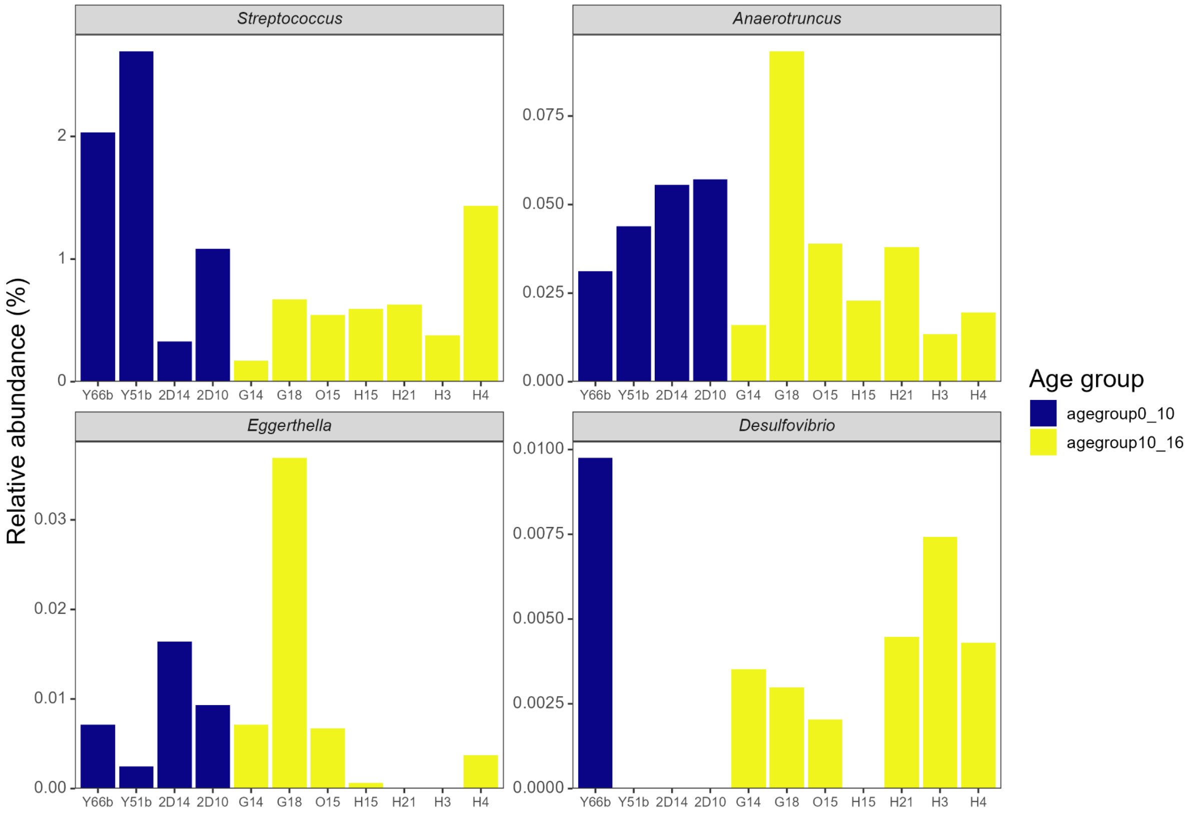
Streptococcus
Anaerotruncus
Eggerthella
Desulfovibrio
Relative abundance (%)
Y66b Y51b 2D14 2D10 G14 G18 O15 H15 H21 H3 H4
Age group
agegroup0_10
agegroup10_16

c

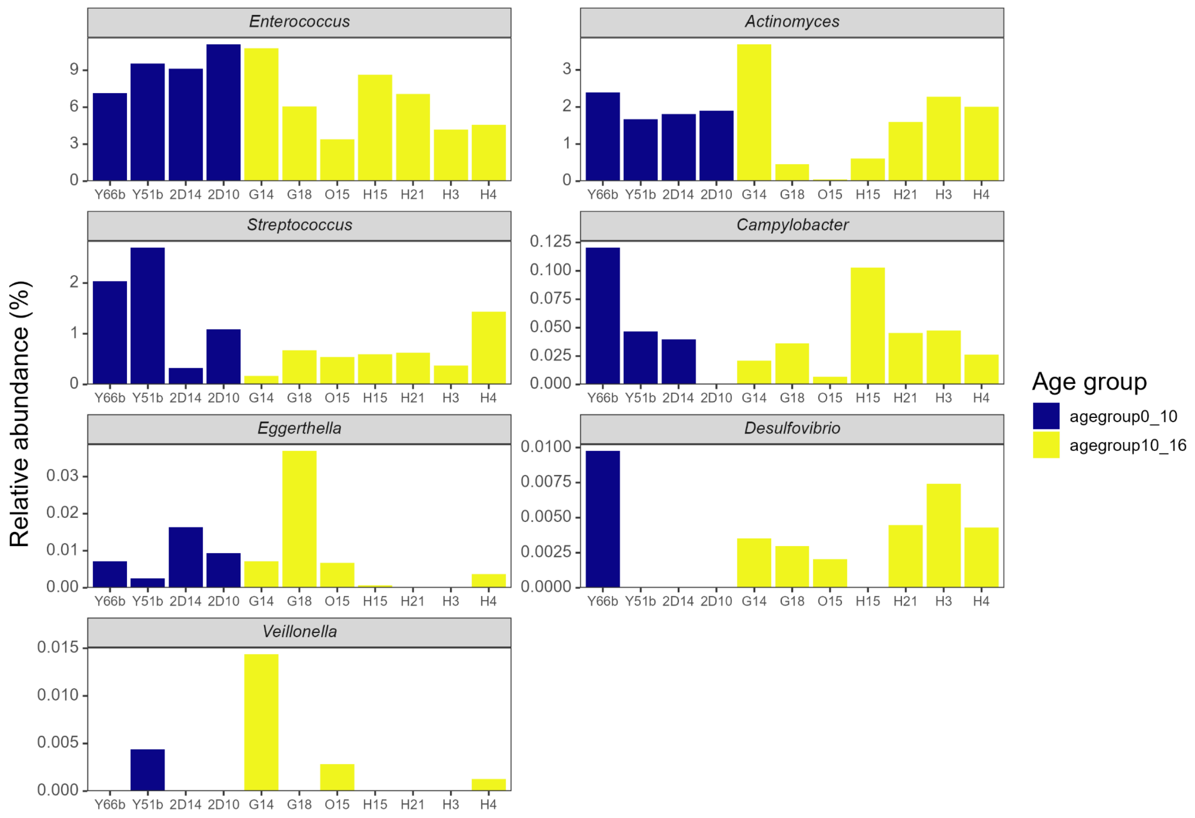

Enterococcus
Actinomyces
Streptococcus
Campylobacter
Eggerthella
Desulfovibrio
Veillonella
Relative abundance (%)
Y66b Y51b 2D14 2D10 G14 G18 O15 H15 H21 H3 H4
Age group
agegroup0_10
agegroup10_16


d

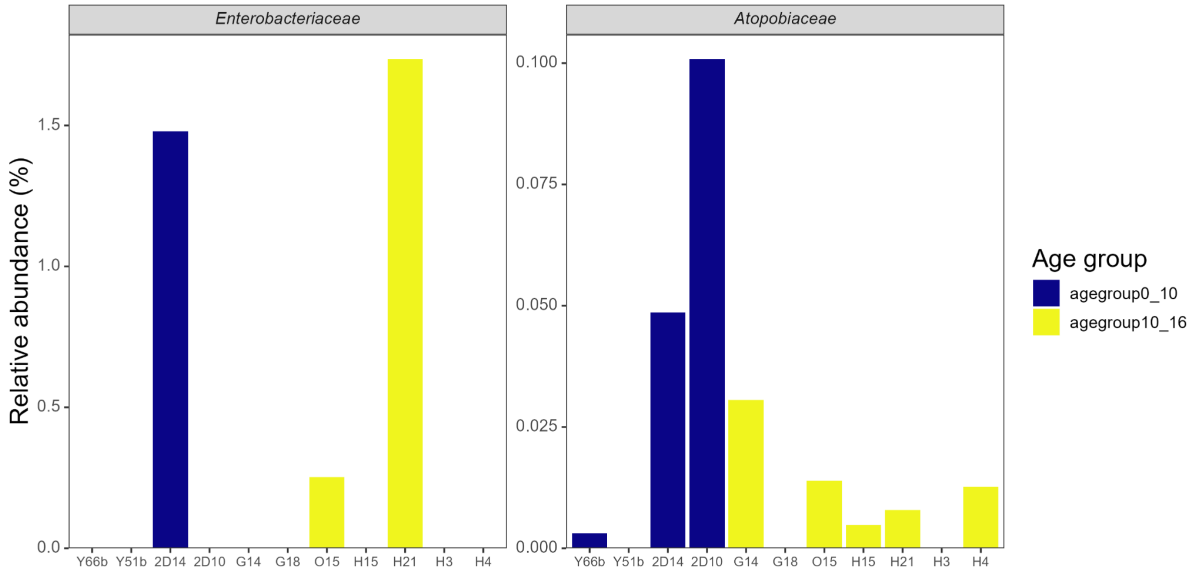

Enterobacteriaceae
Atopobiaceae
Relative abundance (%)
Y66b Y51b 2D14 2D10 G14 G18 O15 H15 H21 H3 H4
Age group
agegroup0_10
agegroup10_16

e

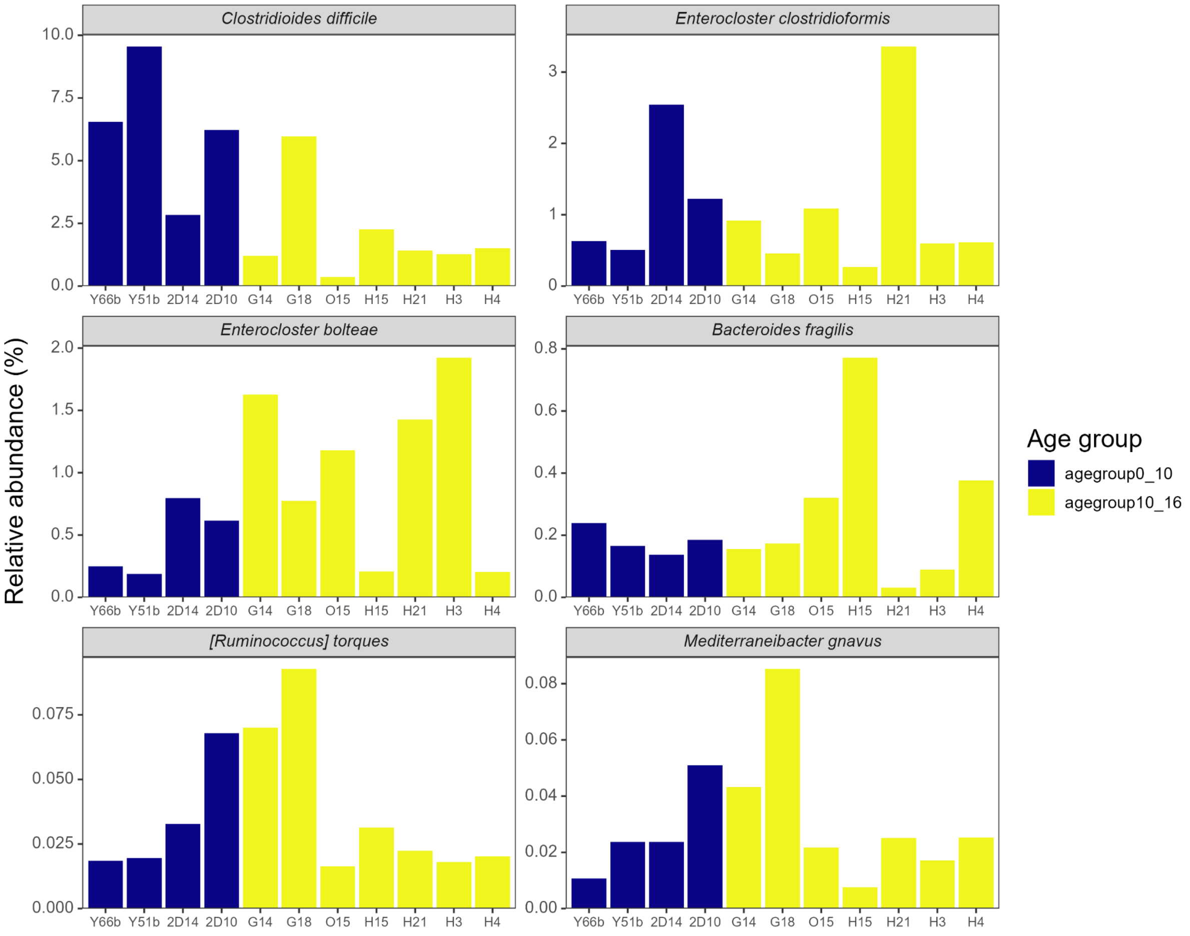

Clostridioides difficile
Enterocloster clostridioformis
Enterocloster bolteae
Bacteroides fragilis
[Ruminococcus] torques
Mediterraneibacter gnavus
Relative abundance (%)
Y66b Y51b 2D14 2D10 G14 G18 O15 H15 H21 H3 H4
Age group
agegroup0_10
agegroup10_16


f

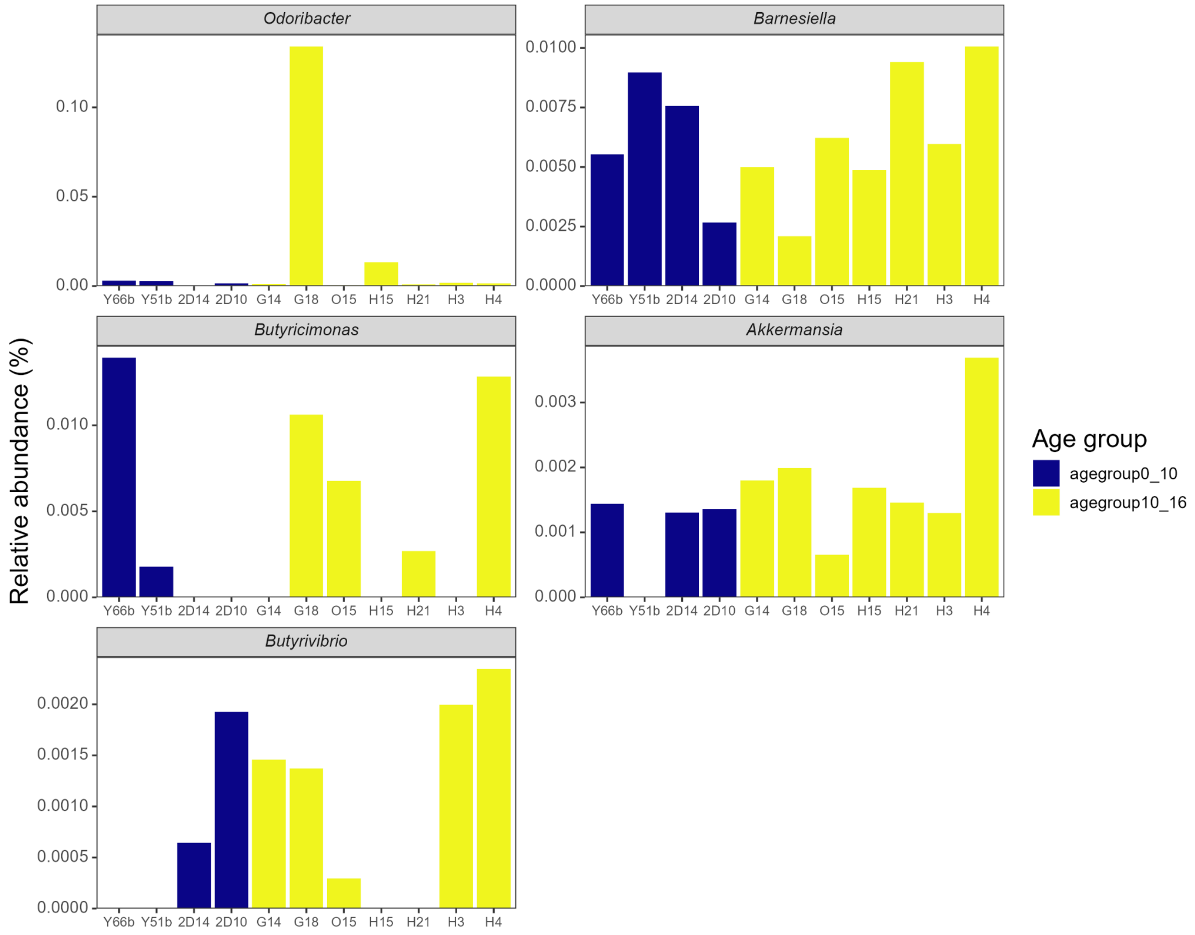
Odoribacter
Barnesiella
Butyricimonas
Akkermansia
Butyrivibrio
Relative abundance (%)
Y66b Y51b 2D14 2D10 G14 G18 O15 H15 H21 H3 H4
0.00
0.05
0.10
0.0000
0.0025
0.0050
0.0075
0.0100
0.000
0.005
0.010
0.000
0.001
0.002
0.003
0.0000
0.0005
0.0010
0.0015
0.0020
Age group
agegroup0_10
agegroup10_16

g

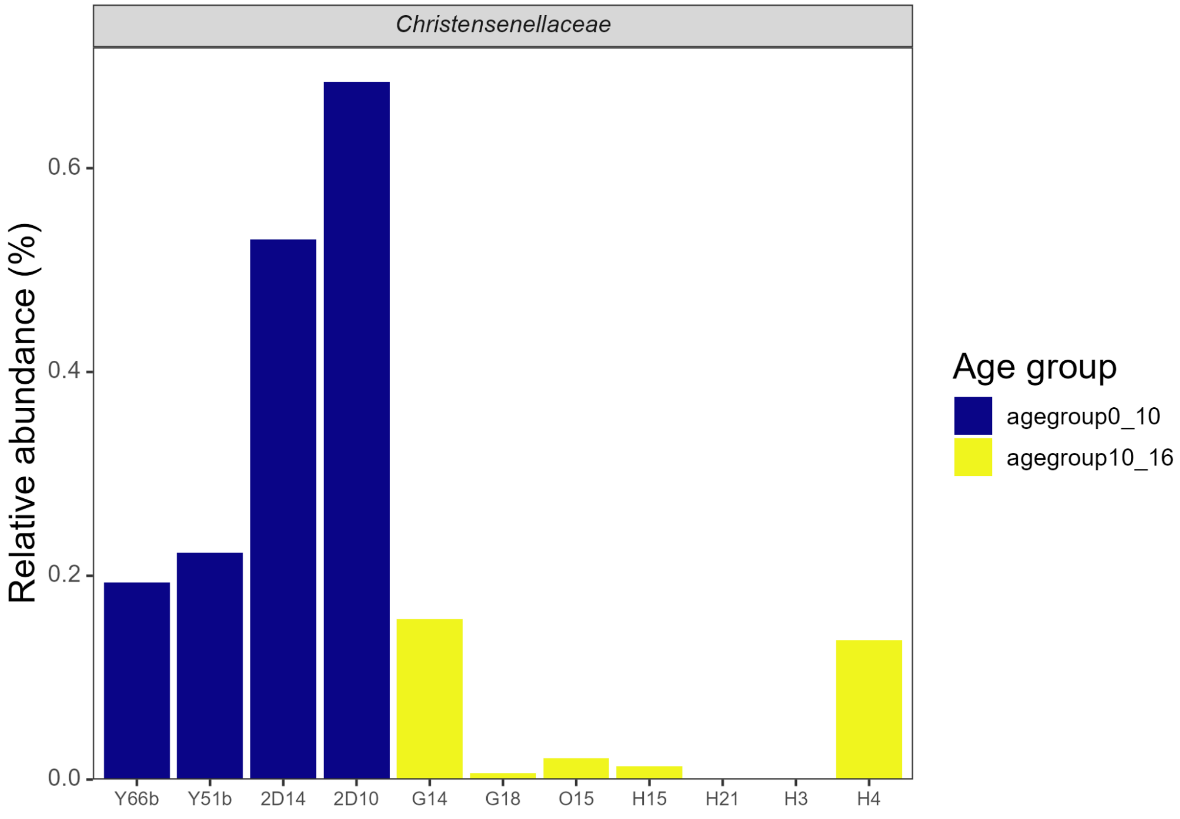


**Figure S9 | Relative abundance of Group 1, 2, and 3 members described in Ghosh et al. (Ghosh et al. 2022) from whole metagenome sequencing | a, Abundances of group 1 genera (decreased with age and associated with healthy aging in humans). b, Abundances of group 2 genera (increased with age). c, Abundances of group 2 genera (associated with unhealthy aging). d, Abundances of group 2 families (associated with unhealthy aging). e, Abundances of group 2 species (associated with unhealthy aging). f, Abundances of group 3 genera (increased with age and associated with healthy aging). g, Abundances of group 3 families (increased with age and associated with healthy aging).**

Figure S10:

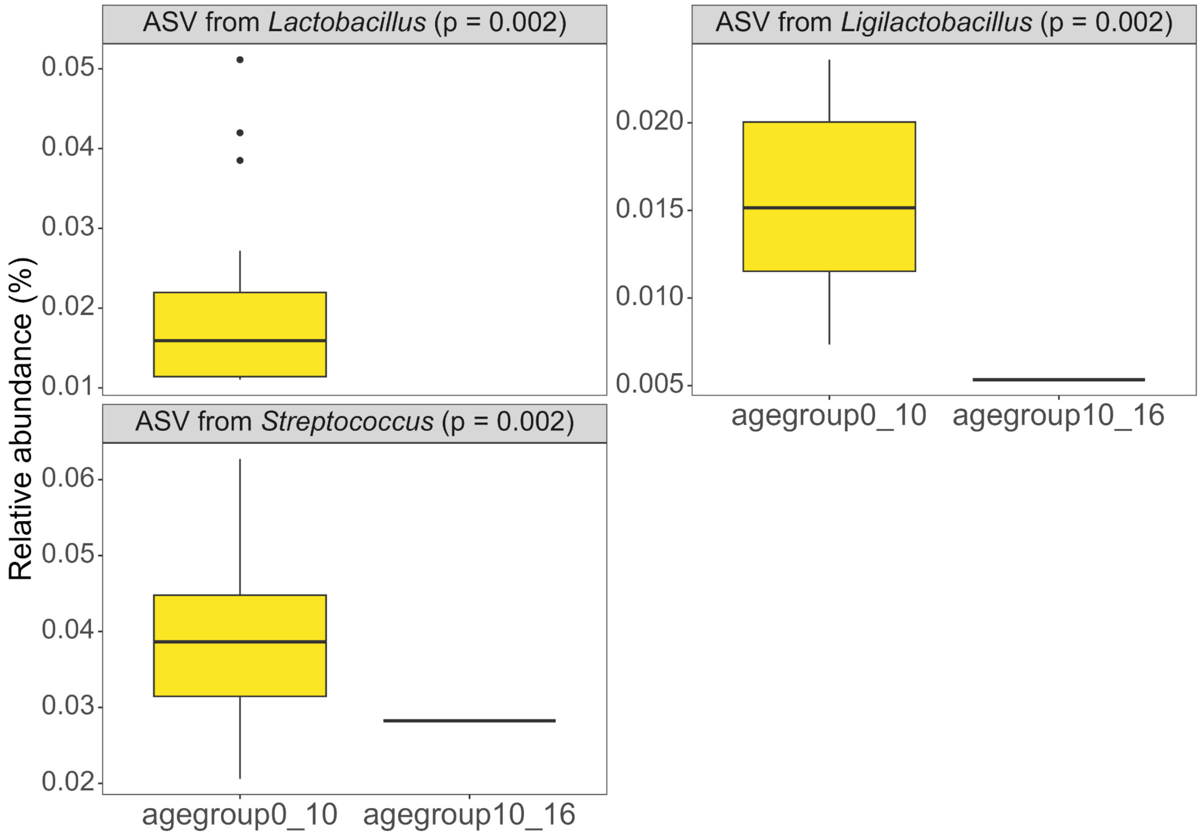


**Figure S10 | Only three ASVs were differentially abundant in young individuals (16S rRNA gene sequencing data).**

Figure S11:

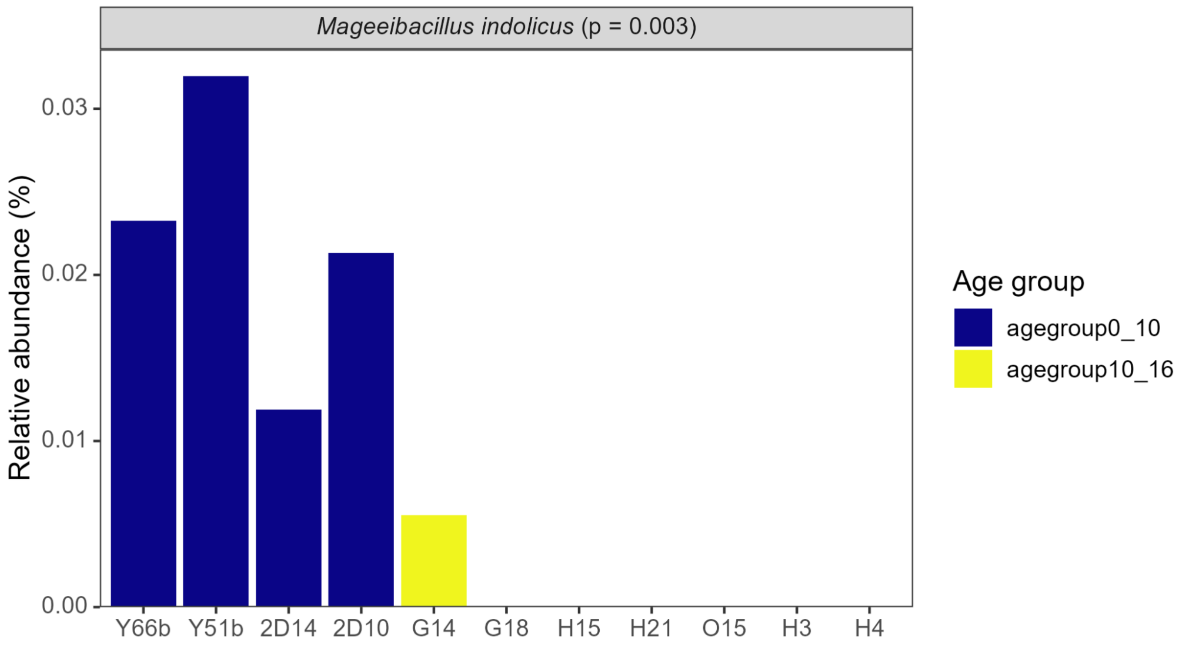


**Figure S11 | *Mageeibacillus indolicus* was significantly more abundant in young naked mole-rats in the whole metagenome sequencing data.**
